# Unraveling the proteome landscape of mouse hematopoietic stem and progenitor compartment with high sensitivity low-input proteomics

**DOI:** 10.1101/2024.05.03.592307

**Authors:** Nil Üresin, Valdemaras Petrosius, Pedro Aragon-Fernandez, Benjamin Furtwängler, Erwin M. Schoof, Bo T. Porse

**Author notes:** These authors contributed equally (Nil Üresin, Valdemaras Petrosius). These authors jointly and equally directed this work (Erwin M. Schoof, Bo T. Porse).

## Abstract

Proteins play a key role in defining cellular phenotypes, yet comprehensive proteomic analysis often requires substantial input material, posing challenges in studying rare populations in complex cell systems. Here, we present an accessible, label-free low-input proteomics workflow that allows for comprehensive proteome coverage reminiscent of classical bulk samples from only 500 cells and showcase its application in murine hematopoiesis. With this approach, we construct a proteomic map of hematopoietic stem and progenitor cell (HSPC) populations isolated by fluorescence-activated cell sorting (FACS) from the bone marrow of a single mouse, identifying approximately 7,000 proteins per cell population. Our study recapitulates the differentiation trajectories along the megakaryocytic-erythroid and granulocytic-monocytic lineages. We specifically focus on the dynamics of transcriptional regulators and provide insights into both known and novel population-specific factors. Furthermore, we extend our exploration to the most primitive stem and progenitor compartment, and identify ADP-Ribosyltransferase ART4 (CD297) as a novel cell surface marker that can potentially be used to enrich for long-term hematopoietic stem cells (LT-HSC). The low-input proteomics workflow presented here holds promise for overcoming the challenges associated with analyzing proteomes of rare cell populations, thereby paving the way for broader applications in biomedical research.

## Introduction

Hematopoiesis, the intricate process of blood cell formation, encompasses a series of differentiation events that originate from multipotent hematopoietic stem cells (HSC) ^1^. These cells progress through various intermediate progenitor stages before ultimately giving rise to mature blood cells, thus ensuring the continuous production of blood throughout an organism’s lifespan. In the murine bone marrow (BM), hematopoietic stem and progenitor cells (HSPCs) are characterized by their expression of the c-kit receptor tyrosine kinase ^2^. The hierarchical model of hematopoiesis traditionally suggests that HSCs serve as the foundation, giving rise to multipotent progenitors (MPP) capable of further differentiating into oligopotent common myeloid (CMP) and lymphoid progenitors (CLP) ^3–5^. HSCs and MPPs reside within the small lineage (Lin)−Sca-1+c-Kit+ (LSK) compartment in murine BM and can be distinguished based on their ability to support long-term engraftment in transplantation assays ^6^. The phenotypic definition of HSC and MPP populations varies, but the widely employed strategy involves utilizing LSKCD150+CD48-CD34-markers to identify dormant long-term (LT) HSCs, while MPP1, characterized by the gain of CD34 expression (LSKCD150+CD48-CD34+), represents a cycling subset of HSCs ^7–9^. MPPs constitute a heterogeneous pool of multipotent cells that exhibit a lineage bias toward either myeloid, lymphoid, or megakaryocyte/erythroid lineages ^10,11^. MPP1-4 populations do not follow a linear developmental trajectory but have been shown to coexist within a pool of multipotent cells displaying different kinetics towards distinct blood cell lineages. Specifically, MPP2 tends to generate megakaryocytic progenitors, MPP3 gives rise to granulocyte-monocyte progenitors, and MPP4 is associated with the lymphoid lineage.

In accordance with the classical tree-like differentiation, MPPs differentiate into CMPs, which subsequently branch into two distinct lineages: granulocyte-monocyte progenitors (GMPs) and megakaryocyte-erythroid progenitors (MEPs) ^3^. To further refine the phenotypic classification as well as the functional characteristics of myeloid cells, additional markers have been proposed. The identification of CD150^+^CD41^+^ cells as megakaryocyte progenitors (MkPs) and CD150^+^FcgR^+^ cells as GMPs has contributed to a better characterization of myeloid cell subsets ^12^. Moreover, the expression levels of the CD105 (Endoglin, ENG) and CD150 (SLAM) markers have enabled the subdivision of novel myeloerythroid precursor stages, such as Pre CFU-E, Pre GM, and Pre MegE, providing a broader characterization of myeloerythroid differentiation ^12^.

While genomics and transcriptomics have provided valuable insights into the rare populations of cells higher in the hematopoietic hierarchy, gaining mechanistic insights into their cellular functions necessitates proteomics studies. It is becoming more evident that proteomic landscapes, inferred by mRNA expression as a proxy, are likely to be an inaccurate reflection of the actual protein content of cells. This was previously demonstrated in seminal work conducted on human erythropoiesis, where mRNA and protein levels did not always correlate for key transcriptional regulators ^13^. Moreover, at the very early progenitor stages, this correlation is particularly weak, likely due to low protein translation rates, emphasizing the importance of direct protein measurements ^13,14^. However, progress in global, antibody-independent proteomics technologies has been hindered by the lack of methods capable of analyzing low-input samples using Mass Spectrometry (MS). Until date, comprehensive MS proteomic analyses of the rarest cell types in mouse BM typically required an input of approximately 50,000 cells to achieve high proteome coverage, necessitating a substantial number of mice for such investigations ^15^. Besides the biological heterogeneity introduced by a large number of mice, collecting rare populations for proteomics notoriously requires extensive sample handling. This process can increase data variability, making it challenging to detect subtle alterations in protein abundances. Recent progress in global label-free proteomics led to the identification of nearly 6,000 protein groups from 25,000 sorted cells within the human HSPC compartment ^16^. Despite the significant decrease in cell-input required for comprehensive proteome coverage, collecting sufficient cells from extremely rare and functionally most relevant samples such as the LT-HSCs remains a substantial challenge, especially in a murine setting.

Given these challenges, there is a need for the development of novel, ultra-low-input MS-based proteomics techniques to facilitate the investigation of rare cell populations. Overcoming this technical barrier would enable the elucidation of cellular differentiation at the proteome level. This goes beyond merely complementing the insights gained from transcriptomics and offers the potential to uncover unique insights that might remain hidden at RNA-based analyses. In this context, the present study aims to overcome the limitations caused by the lack of suitable proteomics technologies for studying the rarest cell populations in the mouse BM.

Here, from a single mouse, we isolated 500 cells from all representative populations within the cKit^+^ compartment, along with both granulocytic and erythroid trajectories, referred to as the “one mouse hierarchy”. Furthermore, from the BM of two pooled mice, we isolated the LT-HSCs and MPP1-4 populations which allowed an in-depth exploration of the protein landscapes defining the most immature HSPC compartment. Subsequently, all samples were subjected to label-free proteomics based on data-independent acquisition (DIA)-MS (WISH-DIA)^17^ on the Orbitrap Eclipse Tribrid MS instrument combined with EvoSep One high-throughput chromatography. In contrast to multiplexed proteomics approaches, our workflow does not require any peptide labeling and only involves minor sample processing steps, and is therefore easily implementable. Remarkably, we identified around 6,500 protein groups per cell type and successfully recapitulated differentiation trajectories in murine hematopoiesis. The resulting data enabled us to reproduce key transcription factor (TF) trends which were previously only measurable in a targeted proteomics setting ^13^. Through our efforts, we aim to contribute to a more comprehensive understanding of early hematopoiesis and pave the way for ultra-low-input proteomics studies in the field of blood cell research and beyond.

## Results

Our initial goal was to generate a comprehensive proteomics atlas covering a wide spectrum of HSPCs, ranging from cells within the LSK compartment toward committed progenitors in both the granulocytic and erythroid lineages **(Figure S1)**. Accordingly, we derived primary cells from the BM of a single mouse and isolated discrete populations based on specific cell surface markers with the use of fluorescent activated cell sorting (FACS) **(Figure S2)**. The cells were further processed with our low input tailored MS-based proteomics workflow to profile the proteomes of the unique populations derived from the BM of a single mouse (**Figure 1a**). From the isolated 500 cells we identified and quantified approximately 7,000 protein groups in each cell type (**Figure 1b**), covering a diverse range of protein classes including plasma membrane proteins and transcriptional regulators. Clustering the samples based on their measured proteomes successfully recapitulated the two main differentiation trajectories, ie. along the granulocyte and erythroid branches (**Figure 1c**). Within the erythroid branch, true MEP should be developmentally upstream of subsequent progenitors in the megakaryocytic-erythroid lineage, yet our analysis positioned them more downstream. This observation is explained by our use of two different antibody panels, each staining for different (sub-)populations. Specifically, the MEP population was isolated in the first panel by the classical definition (CD34-FcgR-), while the remaining erythroid progenitors (ie. Pre CFU-E, CFU-E and Pro Ery) were isolated using the second panel. These results suggest there to be a high level of heterogeneity within the classical MEP gate (CD34-FcgR-), which consequently contains a substantial number of committed erythroid progenitors. The second panel more precisely isolates the more downstream progenitors (ie. Pre CFU-E, CFU-E and Pro Ery), which therefore appear in the correct order in our PCA clustering. Combined, these results underscores the challenge of isolating truly bipotent progenitors through conventional FACS gating, and the utility of MS proteomics data for pinpointing impure cell populations.

**Figure 1.**
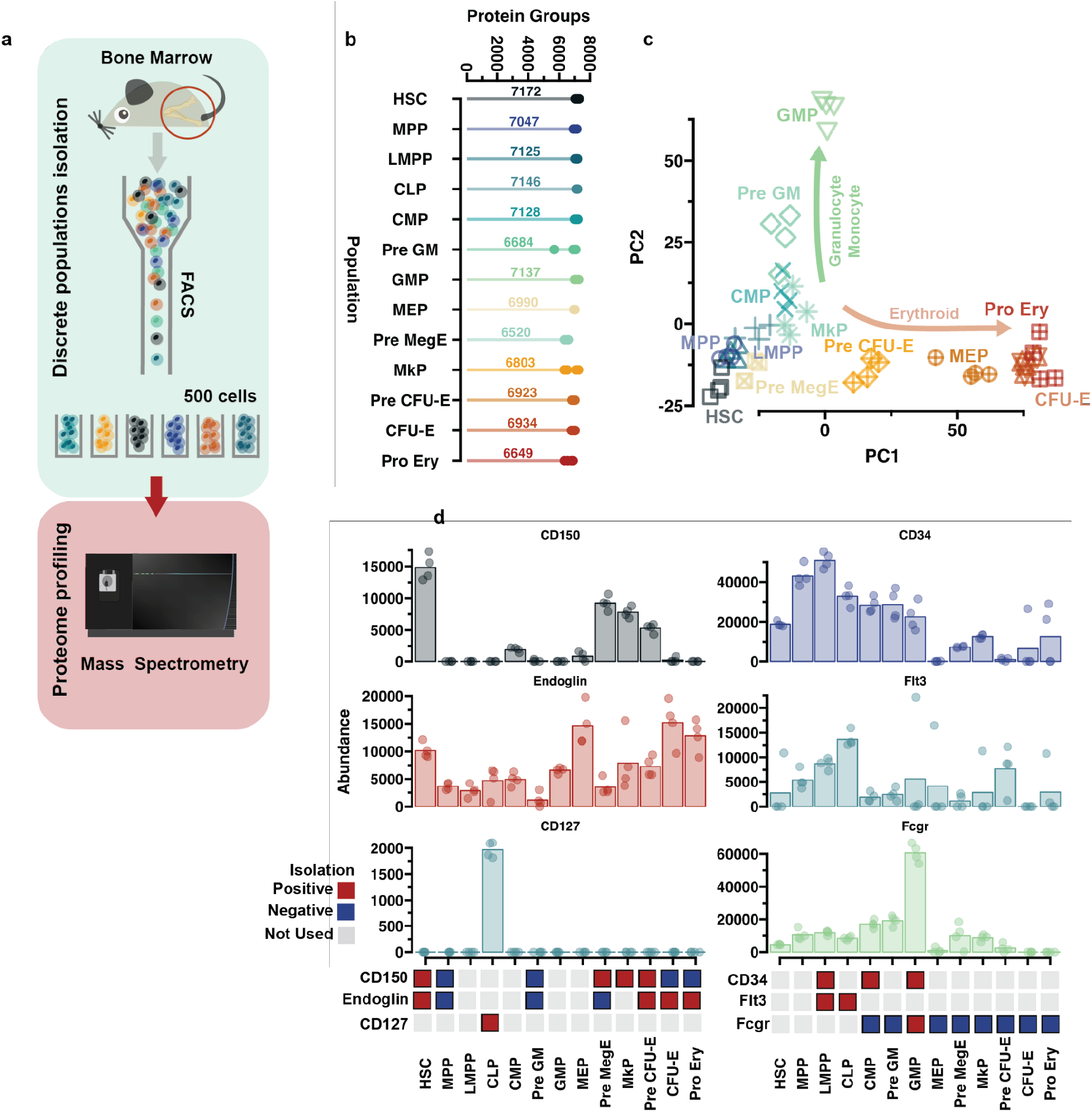
Overview of one mouse hierarchy proteomics data. **a)** Schematic of the experimental workflow. Primary hematopoietic cells were extracted from mouse BM and specific discrete populations were isolated with FACS. Cells were deposited directly into 384 well-plates, where they were lysed and digested. The proteome profiles were measured with the use of MS-based proteomics **b)** Proteome coverage of the analyzed populations. Points represent individual measurements and number represents the average protein group number for the population. **c)** Principal component analysis (PCA) plot of the population. The first two principal components are shown. The specific population is indicated by text and the colors indicate the compartment. Arrows note two emerging differentiation trajectories. **d)** Barplot showing the expression of the cell surface (CD) markers that were used for cell isolation. All the used markers are shown for the populations, however if the markers have been used for isolation (selection) is indicated below the plot by color coding. Bar height represents the average relative abundance of the CD marker and dots note individual measurements.

Next, we set out to further validate the quality of our proteomics data by comparing the expression of cell surface proteins assessed by proteomics and FACS respectively (**Figure 1d**). Here, CD150 exhibited high levels within the HSC population and was undetectable in the MPP population. Additionally, Endoglin expression was highest in the CFU-E and Pro Ery populations, indicating that those markers used for FACS isolation, when quantified in the MS instrument, aligns with the well-established immunophenotype of these cell populations.

### Exploration of transcription factor expression patterns across murine HSPCs

Next, we examined the pattern of TF expression across both granulocytic and erythroid lineages in the ‘one mouse hierarchy’ dataset. We identified specific expression of master regulators, such as SPI1 (PU.1) and CEBPA in the granulocytic lineage ^18–20^, and TAL1 and GATA1 in the erythroid trajectories (Figure 2a) ^21,22^. CEBPA is highly specific for the myeloid lineage, with its expression progressively increasing from CMP towards pre-GM and reaching the highest level in GMP (Figure 2a). To potentially capture novel TFs involved in early hematopoietic differentiation, affinity propagation (AP) clustering was applied on the 380 TFs that were identified in our data (obtained from the animalTFDB 4.0). AP finds the optimal number of representative examples in the data and clusters all the transcriptional regulators based on their similarity to the examples, **(see Methods)**, highlighting unique patterns of TF expression in the megakaryocytic-erythroid and granulocytic-monocytic lineages (Figure 2b-c). Through this analysis, we both validated known, and uncovered novel TF expression patterns associated with different stages of differentiation, highlighting the diversity in TF regulation throughout differentiation.

**Figure 2.**
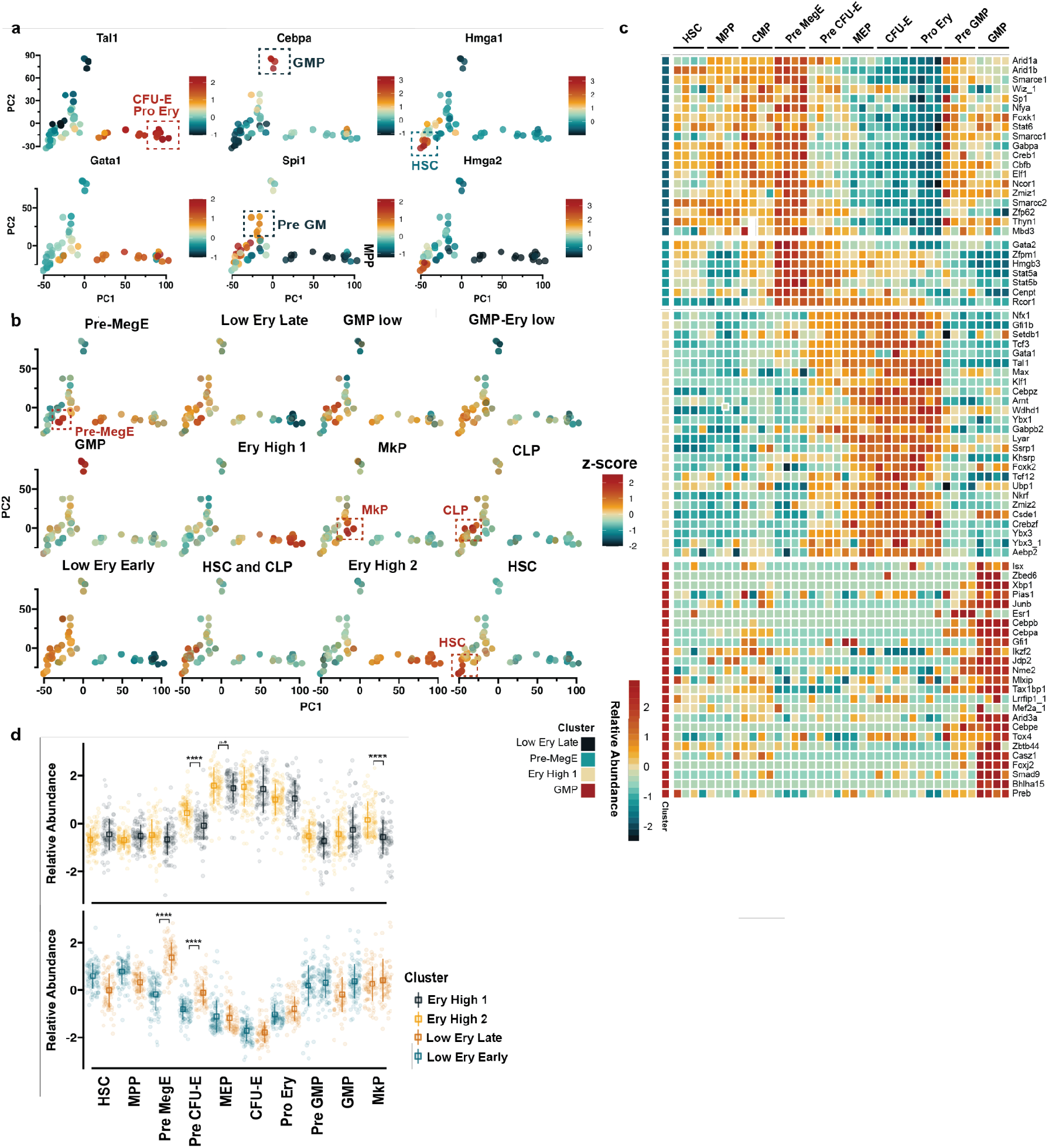
Identification of transcription factor expression trends. **a)** Trends of selected transcription factors. PC1 and PC2 are used for the x and y axis. The color indicates the relative TF abundance. Factors involved in erythroid, granulocyte and HSC function were selected based on literature. Some distinct populations are indicated by rectangles and text **b)** Affinity propagation (AP) clustering results of TF expression. All the TFs from an AP cluster are overlaid, with the color intensity corresponding to normalized abundance. If the overall trends match, it results in a bright color, while if they do not, the different colors mix and no clear pattern can be observed. **c)** Heatmap of selected clusters of transcription factors **d)** Scatter plots showing the transcription factor abundance distribution in different populations highlighting the major difference between the erythroid clusters. The top and bottom sections show TF abundances that are either increasing or decreasing, respectively, for the erythroid populations. The box shape indicates the mean relative abundance and the line represents the standard deviation. KS test was used to determine significant differences. n.s. - not significant, **** p-value < 0.0001. Full cluster overview is available in the appendix.

The AP clustering analysis revealed population-specific TFs as well as shifts in TF expression patterns across different trajectories. The high mobility group proteins, HMGA1 and HMGA2, revealed high expression in HSCs (Figure 2a-b). Previous research has linked HMGA2 to the self-renewal potential of fetal HSCs, indicating an important role for this chromatin modulator in HSC function ^23^. We next focused on the dynamics of TF abundances in erythroid differentiation, and observed a shift in their expression pattern. A cluster of TFs exhibited low expression in the erythroid progenitor hierarchy during CFU-E and Pro-Ery stages (“Low Ery Late”), while another set of TFs showed low expression starting earlier in the trajectory at the PreCFU-E stage (“Low Ery Early”). Similarly, we identified clusters of TFs with increased expression during the erythroid trajectory, referred to as “Ery High 1” and “Ery High 2”, the latter starting earlier in the PreCFU-E stage as well as exhibiting higher abundance in the MkP population (Figure 2b**,d**). GMPs exhibited a distinct cluster of TFs, including both well-established and previously undocumented factors. Among the known factors, different members of the CEBP family, CEBPA, CEBPB, and CEBPE, showed the highest protein levels detected in GMPs (Figure 2c). Furthermore, JUNB, known to be a crucial transcriptional regulator in myelopoiesis, demonstrated highest abundance in the GMP cluster ^24^. Next, we conducted a comparative analysis of transcriptome-proteome dynamics during differentiation based on data extracted from the BloodSpot database ^25^ and the data generated in the present work **(Figure S3)**. This comparison illustrates a good correlation between the transcript and protein levels of the CEBPA, however, the relationship for CEBPB and CEBPZ appears to be more complex **(Figure S3a)**. Our data also points towards GMP-specific transcriptional regulators that have not been previously described in the context of granulocytic differentiation, including CASZ1, FOXJ2, and PREB, all demonstrating high levels in GMPs, where the correlation between mRNA and protein levels are less pronounced. This is exemplified by FOXJ2, a member of the Forkhead Box TFs, which exhibits high transcript levels in HSCs that gradually decrease during differentiation **(Figure S3b)**. In contrast, our proteome data identifies FOXJ2 as a highly specific TF for GMPs, demonstrating the importance of using protein level data.

To further assess the quality of our generated data, we compared TF expression along the erythroid differentiation trajectory to data obtained with targeted proteomics techniques from human samples ^13^. Out of the 84 orthologs included in the original work, 80 were quantified in our dataset and were in alignment with previously reported findings^13^. Although the expression patterns cannot be directly compared due to the difference in species, we observed highly complementary TF abundance dynamics along the erythroid differentiation trajectory. This is exemplified by the decreasing abundance of SPI1 and FLI1, while levels of KLF1-3 and TAL1 were increasing **(Figure S4)**, underlining that our proteomics data can be used to identify novel TF expression patterns in the isolated populations.

### Identification of population-specific cell surface markers

Next, we explored cell surface proteins specific to different HSPC populations, considering their potential for immunophenotypic isolation, either in combination with or as alternatives to existing isolation schemes. Utilizing the MS-derived Cell Surface Protein Atlas (CSPA) as a reference ^26,27^, we investigated the abundance of 323 surface marker profiles (those that were detected in our data, see methods) across different cell populations (Figure 3a). First, we explored candidate cell surface markers specific to the HSC population within the one mouse hierarchy dataset. This population was isolated based on the expression of CD150 and ENG, and accordingly, we identified SLAMF1 and ENG as surface proteins upregulated in HSCs compared to the LSK population (Figure 3b**,c**). Conversely, the cell surface proteins CD48 and CD34 exhibited downregulation in the HSC population (Figure 3d). These surface markers served as clear reference points supporting the validity of our proteomics data. Next, we investigated potential HSC-specific markers. Notably, ADP-ribosyltransferase ART4 (CD297) emerged as a candidate marker showing specificity for HSCs (Figure 3e). ADP-ribosylation is a post-translational modification that plays a role in various cellular processes, including the regulation of glucose metabolism ^28^. Elevated levels of ART4 might be linked to the precise control of metabolism in HSCs. Furthermore, CLEC2D, a C-type lectin, is upregulated in HSCs. CLEC2 family proteins exhibit a diverse range of functions, including mediating cell-cell adhesion. Cell adhesion molecules play an active role in the signaling cascades that regulate HSC quiescence and self-renewal ^29^. These two surface proteins are likely candidates for enhanced purification of HSCs.

**Figure 3.**
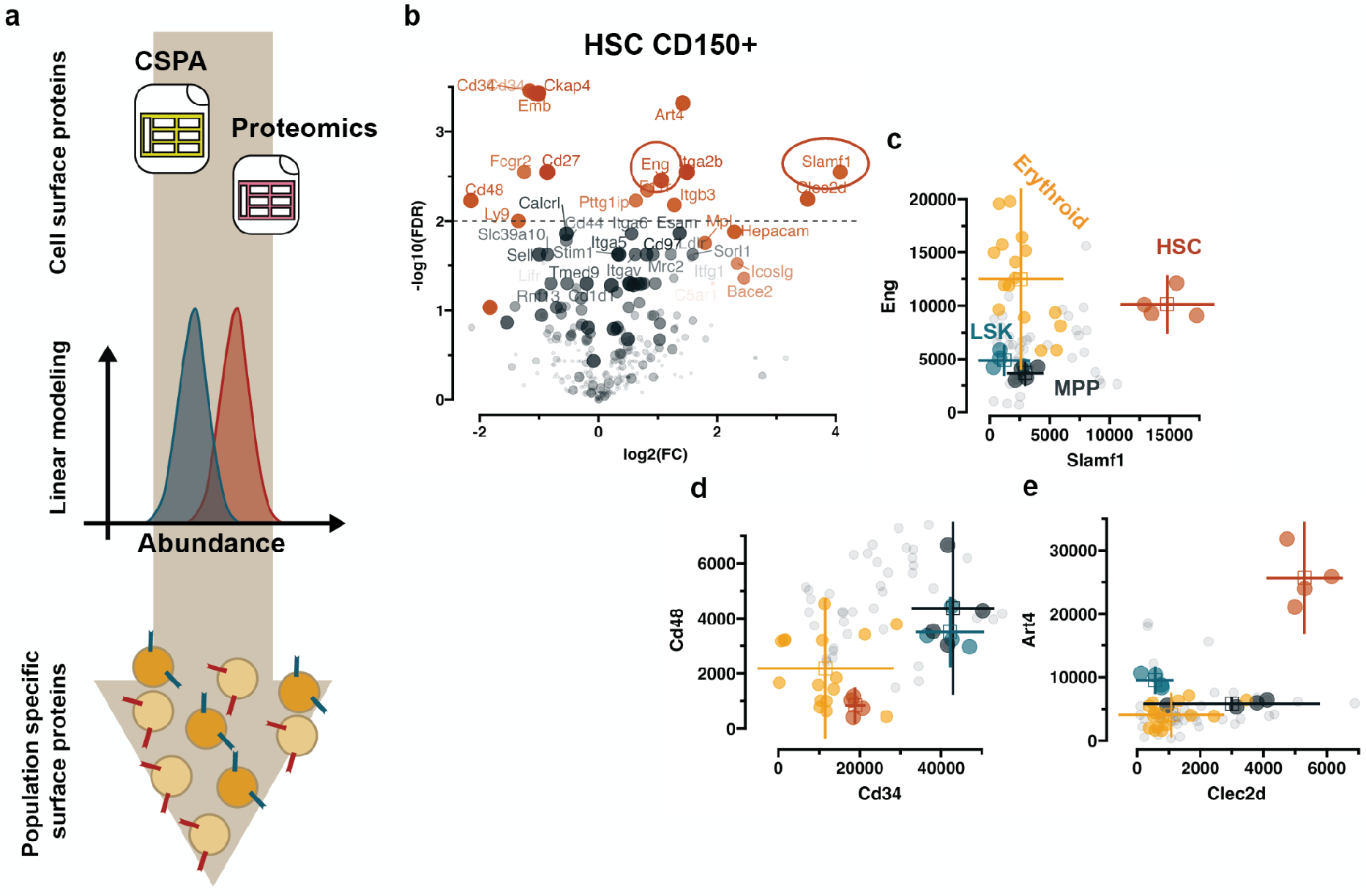
Inference of population specific cell surface markers. **a)** Schematic of workflow for marker identification. CSPA: Cell Surface Protein Atlas **b)** Volcano plots showing potential the most abundant proteins for HSC CD150+ populations, relative to MPP CD150-. The log2 fold-change (FC) is plotted on the x-axis and the -log10 false discovery rate (FDR) on the y. Dot size indicated the mutual information value. Proteins that would be considered hits are color coded. **c-e)** Scatter plot showing protein abundances of specific proteins noted on the axis. Endoglin (ENG) and CD150 (SLAMF1) are shown as a reference to show that the workflow can identify population specific cell surface proteins. The Erythroid populations are color coded separately as they are also positive for ENG. CD34 and CD48 are shown as further evidence. Finally, ART4 and CLEC2D are shown as potentially novel CSP that could be used to isolate the same cell population. The lines indicate 95% confidence interval (std * 1.96) for each protein. Grey points note all the remaining populations that are not otherwise marked.

Furthermore, the cluster of differentiation markers CD180, CD68, and CD93 were identified as specific to CLPs and can be tested either in combination or as alternatives to the IL7 receptor for the isolation of CLPs (Figure 4a**,b**). Additionally, we pinpoint cell surface proteins specific to GMPs, such as EMB, which, similar to FcgR, belongs to the immunoglobulin superfamily and could serve as a potential cell surface marker for the isolation of GMPs (Figure 4c**,d**). In both mouse and human BM, MEPs are purified based on the absence of specific markers, resulting in a heterogeneous population. It is likely that through the discovery of surface markers that are specific to the biological, bipotent MEP population, more pure cell populations can be obtained. Our data suggests Pre-MegE cells to be positioned upstream of not only MEP, but also both MkP and Pre-CFU-E, suggesting that the immunophenotypic pre-MegE population better represents bipotent MEPs (Figure 1c). Our analysis points towards cell surface proteins such as CD47 and ADAM10 that are enriched in the Pre-MegE population and may potentially be used for further resolving this population (Figure 4e**,f**). Overall, these surface markers offer the potential to improve current FACS isolation strategies and can be used in functional assays to evaluate the lineage output of sorted populations.

**Figure 4.**
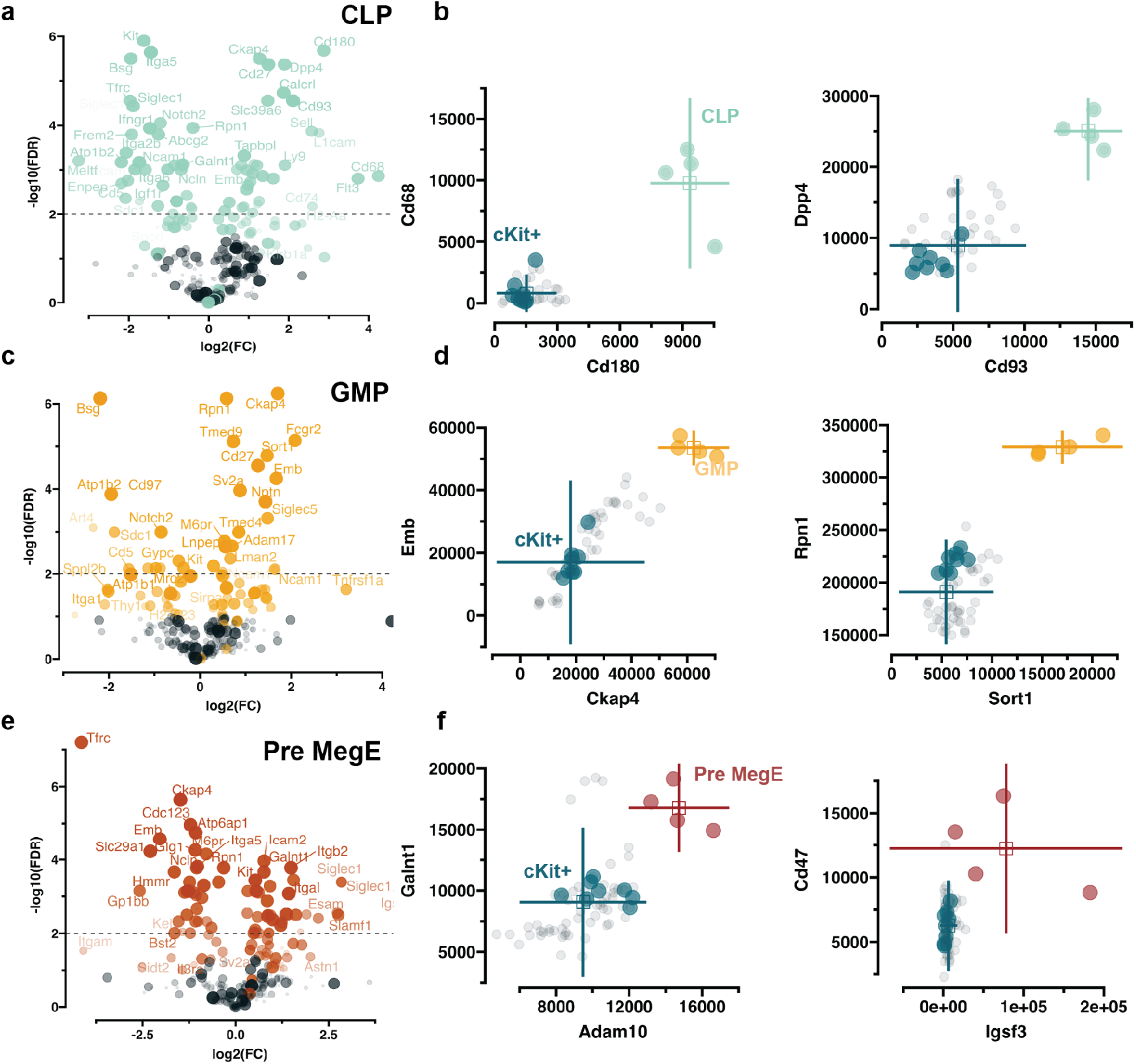
Identified cell surface markers for progenitor populations. **a, c, e)** Volcano plots showing potential the most abundant proteins for CLP, GMP and Pre MegE populations relative to cKit+. The log2 fold-change (FC) is plotted on the x-axis and the -log10 false discovery rate (FDR) on the y. Dot size indicated the mutual information value. Proteins that would be considered hits are color coded. **b, d, f)**

### Proteomic profiling of the LT-HSC and MPP populations

Having demonstrated the robustness of our low-input proteomics workflow, we proceeded to investigate the early stem cell compartment. The most comprehensive proteomics investigation to date focused only on LT-HSCs and MPP1 and required 400,000 cells for input, as well as peptide labeling and fractionation ^9^. Our goal was to generate a proteomics dataset covering LT-HSCs and MPP1-4 from merely 500 cells, and with minimum sample handling to enable such studies at a routine level. Due to the existing limitation of BD cell sorters, which prevents the simultaneous isolation of cell types (e.g. 4-way sort) into 384-well plates, and the rare nature of the populations under study, we opted instead to pool BMs from two mice. This approach allowed us to sort LT-HSCs and MPP1-4 populations from the same source of cells, facilitating a comprehensive analysis of these early stem and progenitor cells (Figure 5a**, Figure S2 Panel 3)**. Despite the primitive nature of these stem cell populations, we successfully quantified approximately 5,500 proteins per population (Figure 5b). The PCA analysis demonstrated a cohesive clustering among all the studied populations, providing further support to the robustness of our workflow (Figure 5c). LT-HSCs demonstrated an enrichment of proteins associated with metabolic processes, including iron homeostasis and proteostasis, highlighting the importance of metabolic regulation in LT-HSCs. The ferroxidase FTH1, guanine deaminase GDA that is involved in purine metabolism, and regulators of proteostasis such as SAMHD1 and TGM2, were upregulated in LT-HSCs. Additionally, high levels of the linker histone H1F0 in LT-HSCs suggests a compact chromatin structure, characteristic of these quiescent stem cells. On the other hand, MPP1 exhibited an enrichment of proteins involved in cell-cycle regulation, mitotic progression, and spindle assembly. These include cell-cycle related proteins such as CDK6 and CCNA2, as well as proteins involved in mitotic spindle assembly such as TPX2, KIF22, KIF11, and NUSAP1 in line with initiation of proliferation (Figure 5d). In previous efforts to profile the proteome level differences of HSCs and MPP1, only 47 proteins were identified as differentially expressed at a False Discovery Rate (FDR) of 10%, with none passing the more stringent 1% cut-off ^9^. At 10% FDR, our data revealed 830 proteins as differentially expressed, marking an overall 15-fold increase (Figure 5d).

**Figure 5.**
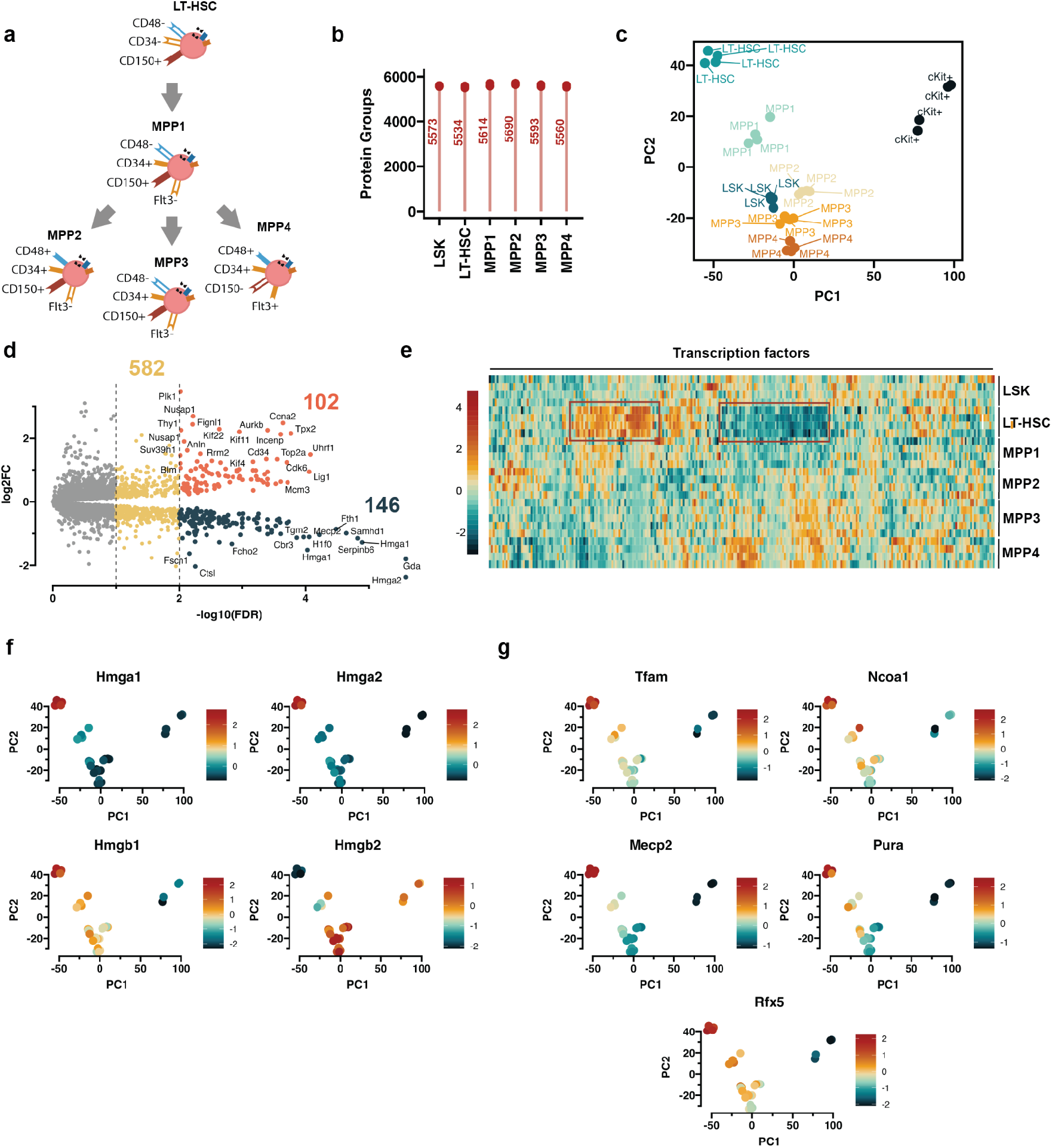
Proteomic profiling of the HSC MPP populations. **a)** Schematic of isolated populations of interest. **b)** Proteome coverage of the analyzed populations. Points represent individual measurements and number represents the average protein group number for the population. **c)** Principal component analysis (PCA) plot of the population. The first two principal components are shown. The specific populations are indicated by text and color. **d)** Volcano plot showing the top hits between HSC and MPP1 populations. Positive log fold-change (logFC) represents increased abundance in MPP1 and decreased if the logFC is negative. Two filtering thresholds are shown 1% and 10%. The number note the number of proteins **e)** Heatmap showing the expression of transcription factors. **f-g)** PCA plots with overlaid TF abundances represented by color. The specific transcription factors are indicated above the plots.

By examining TF abundance levels in the different populations, we observed a distinct pattern of transcriptional regulators specific for LT-HSCs (Figure 5e). Notably, high-mobility group proteins exhibited a unique expression pattern in early stem cells. Specifically, HMGA1 and HMGA2 displayed high specificity in LT-HSCs. This observation aligns with the previous proteomics study, where HMGA1 and HMGA2 were identified among ∼20 proteins exhibiting increased expression in LT-HSCs ^9^. In contrast, while HMGB1 exhibited elevated levels in LT-HSCs, it also showed increased abundance in MPP1-4. Interestingly, HMGB2 appeared to have low expression in LT-HSCs and MPP1 but showed higher levels in MPP2-4 (Figure 5f). High mobility group proteins serve as non-histone chromatin modulators also known as “architectural transcriptional factors” involved in various processes ^30,31^. While HMGA2 and HMGB1 have been shown to play a role in myeloid malignancies as well as other cancers ^32–34^, their temporal involvement in the adult HSPC compartment has not been documented.

Moreover, the expression of Transcription Factor A Mitochondria (TFAM), a protein that binds to mitochondrial DNA and plays a role in transcription, replication, and packaging of mitochondrial DNA ^35^, appears to be specifically associated with LT-HSCs. Its levels decrease as cells transition from the MPP1-4 stages. Similarly, MECP2 (methyl-CpG binding protein 2) exhibits enrichment in LT-HSCs, whereas its abundance decreases in the MPP compartment (Figure 5g). MECP2 is mutated in Rett syndrome ^36^, a rare genetic neurological disorder, but has not previously been reported in hematopoiesis. Indeed, when examining the public gene expression data (Bloodspot database) ^25^, we could not find any indications of upregulation of this gene in LT-HSCs. The enrichment of MECP2 in the most immature stem cells prompts further exploration of its potential role in HSC function. In summary, our data highlights both known and novel transcriptional regulators that appear to exhibit enrichment in LT-HSCs compared to the downstream MPP compartment. However, future investigations are needed to elucidate their functional roles in maintaining stem cell quiescence.

We further leverage the generated data to determine potential cell surface markers that can be used to distinguish LT-HSC and MPP1 with our previous approach (Figure 3). As a quality control measure, we could confirm differential abundance of SLAMF1 and CD34 between LT-HSCs and MPPs (Figure 6a**,b**). Consistent with the known immunophenotype of these populations, LT-HSCs exhibit high levels of SLAMF1 and low levels of CD34. Similar to the broadly isolated CD150+ HSC population in the one mouse hierarchy dataset (Figure 3e), ART4 demonstrated a high specificity for LT-HSCs (Figure 6c). Therefore, our data suggests that the simultaneous use of CD150 and ART4 could potentially enrich for LT-HSCs. Our data further illustrates that THY1 (CD90) serves as a distinguishing factor for identifying LT-HSCs, with LT-HSCs being characterized by low THY1 expression compared to the MPP compartment (Figure 6c). In contrast to its well-established role in human hematopoiesis ^5^, CD90 is not commonly utilized to isolate mouse HSCs. It is interesting to observe that the most frequently employed markers CD34, CD38, and CD90 tend to exhibit divergent patterns of expression between mouse and human Overall, we identify tentative cell surface markers that could be utilized for LT-HSC isolation.

**Figure 6.**
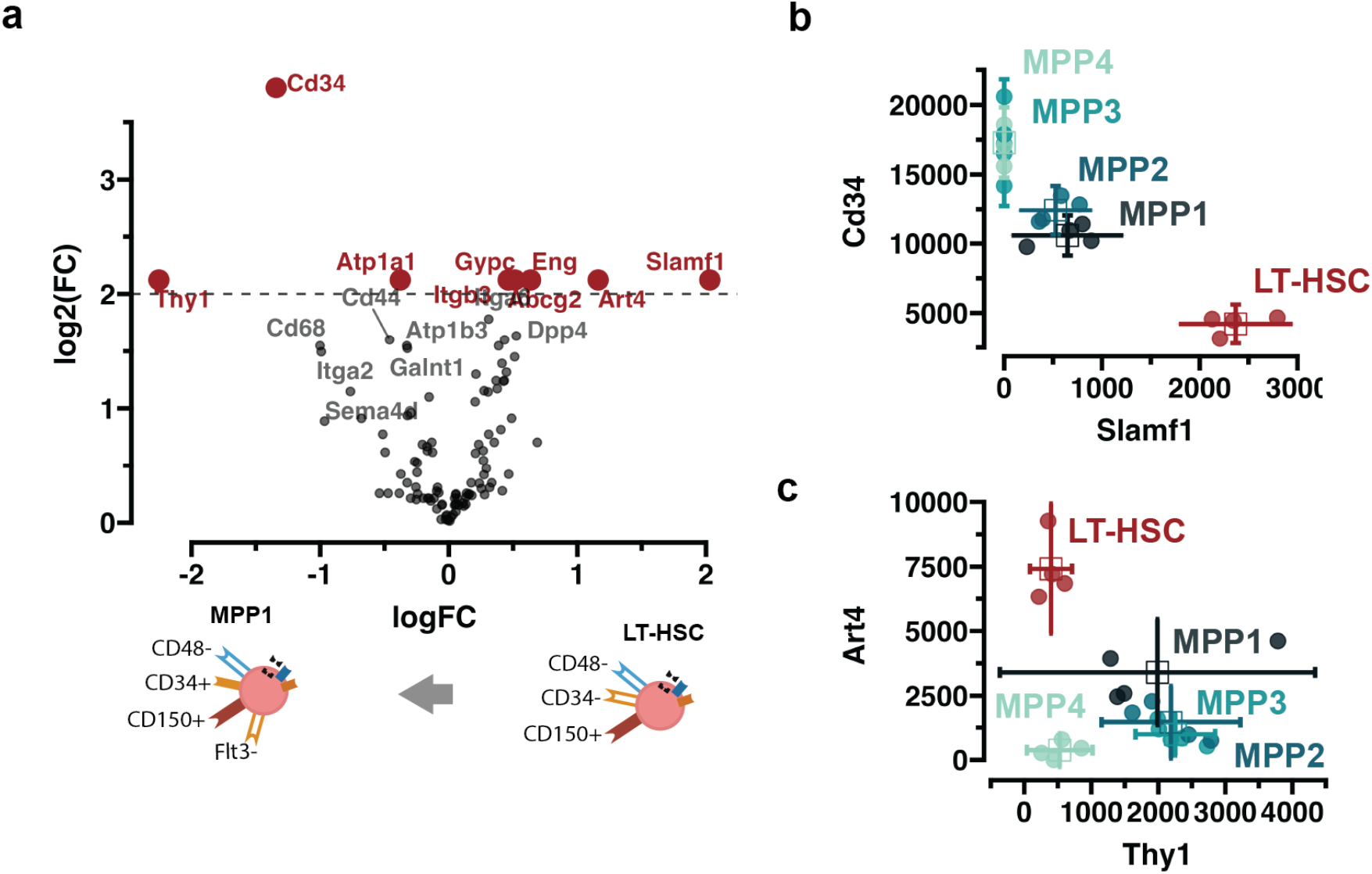
Distinct cell surface markers for LT-HSC population. **a)** Volcano plot of difference in cell surface protein abundance of LT-HSC population relative to MPP1. **b-c)** Scatter plot showing protein abundances of specific proteins noted on the axis. The lines indicate 95% confidence interval (std * 1.96) for each protein. Populations are color coded and also noted in text.

## Discussion

In this study, we showcased the potential of employing ultra-low-input proteomics in a complex biological system. Through our refined protocol, which minimizes sample handling that relies entirely on non-proprietary off-the-shelf consumables, this study is the first to report a comprehensive proteomic map of the hematopoietic hierarchy extracted from the BM of a single mouse. Despite the limited number of cells required, the workflow yielded high proteome coverage and captures expected biological trends underlining the quantitative quality of the generated data.

Latent variables analysis could recapitulate the trajectories of cellular differentiation noting a clear bifurcation from HSC into erythroid and granulocytic lineages from the proteomes profiles (Figure 1c). We capitalized on this trend by applying affinity propagation clustering to discern transcriptional regulator expression patterns in the discrete populations (Figure 2). Expected transcriptional regulators were aggregated together according to their established function, and we identified novel candidates which have not been previously reported to be distinctly regulated at the mRNA level. FOXJ2 appears to be a highly specific TF for GMPs and has not previously been implicated in granulocytic differentiation. We detected nearly 400 proteins classified as TFs, representing 25% of the entire murine ‘TF-ome’. Typically, as low-abundant proteins, measuring TFs relied on targeted proteomics methods such as selected reaction monitoring (SRM) ^13,37^. Here, our sensitivity-tailored DIA workflow provides a high coverage of TFs without the need for synthetic peptides that are typically required for SRM experiments. Importantly, our data highlights transcriptional regulators upregulated in LT-HSCs, including MECP2 and HMGA family members, factors which have not previously been characterized in detail in the context of the adult murine hematopoietic system (Figure 5g). Overall, these findings not only clearly establish that our data paves the way for future functional analyses, but also underscore the importance of direct protein-level measurement.

It is a long-standing holy grail in the field to identify surface markers that can effectively distinguish between LT-HSCs and their nearest progenitors. In the most extensive proteomic dataset to date, the comprehensive examination of LT-HSC and MPP1 revealed only a few surface proteins that are differentially expressed ^9^. In our analysis, ART4 emerged as a candidate LT-HSC specific surface marker. As an ADP-Ribosyltransferase involved in regulating metabolic pathways, ART4 is a likely candidate as an LT-HSC marker, especially given the tight control of energy metabolism in quiescent stem cells.

Finally, while we opted to apply our workflow in a well-described stem cell hierarchy, its potential extends to exploring proteomes of clinically relevant, rare cell types, including circulating tumor cells and cancer stem cells. The possibility to generate comprehensive proteomic profiles with an input of only 500 cells represents a significant advancement, bringing proteomics studies closer to applications in clinical and other biomedical settings where limited sample availability has precluded MS proteomics analysis to date. As a corollary, we also note that our approach has ethical implications, as we have drastically reduced the number of experimental mice required for these types of investigations. With the high-sensitivity MS instruments and intelligent data acquisition strategies, obtaining a robust proteomic dataset from even fewer than 500 cells will soon become feasible. This makes low-input proteomics an attractive workflow for investigating rare, yet biologically relevant cell types.

## Materials and Methods

### Isolation of HSPCs from mouse BM and FACS sorting

Four 14-week-old female C57BL/6 mice were sacrificed by cervical dislocation and leg bones (femur, tibia and iliac crest) and spines were dissected. The bones were crushed three times with 10 mL PBS/3% FBS and the suspension was filtered through a 70μm cell strainer. The filtered suspension was filled up to 50 mL and centrifuged at 300g for 10 min at 4°C. To enrich for cKit^+^ cells, the pellet from each BM sample was resuspended in 392µl PBS/3% FBS and 8µl CD117 (cKit) MicroBeads (Miltenyi Biotec, catalog number 130-091-224) and incubated on ice for 30 mins. Magnetic separation was carried out using LS Columns according to the protocol provided by Miltenyi Biotec. cKit-enriched BM was stained with antibodies for 30 min at 4°C (see panels 1 and 2 below for one mouse hierarchy). The stained cells were washed three times with PBS to remove the residual serum by centrifuging at 300g for 10 min at 4 °C. The pellet was resuspended in PBS containing 1:1000 (v/v) 7-AAD viability dye (Invitrogen, catalog number A1310). 500 cells per well were sorted on a BD FACSAria III sorter using a 100-micron nozzle in 4-way purity precision mode into an Eppendorf twin.tec 384 LoBind plate containing 1µl of 0.2% DDM, 80mM TEAB pH8.5 lysis buffer. For the HSC and MPP compartment, leg bones from two mice were combined (8 mice total, n=4) and stained with panel 3. The cell sort was done using a 100-micron nozzle in a purity mode on the BD FACSymphony S6 cell sorter. After the sort, plates were briefly spun down, snap-frozen on dry ice, and boiled at 95**°**C for 5 min to complete the cell lysis using a Veriti thermal cycler (Applied Biosystems). Subsequently, the heated plates were allowed to cool down on ice and proteins digested by adding 1µl of 10ng Trypsin Gold (Promega, catalog number V5280) in 100mM TEAB and incubating at 37**°**C overnight. The next day, each well was acidified by the addition of 1 μL 4% (v/v) trifluoroacetic acid (TFA). All liquid dispensing into 384-well plates was done using an I-DOT One instrument (Dispendix).

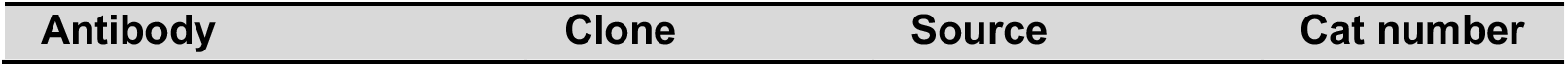
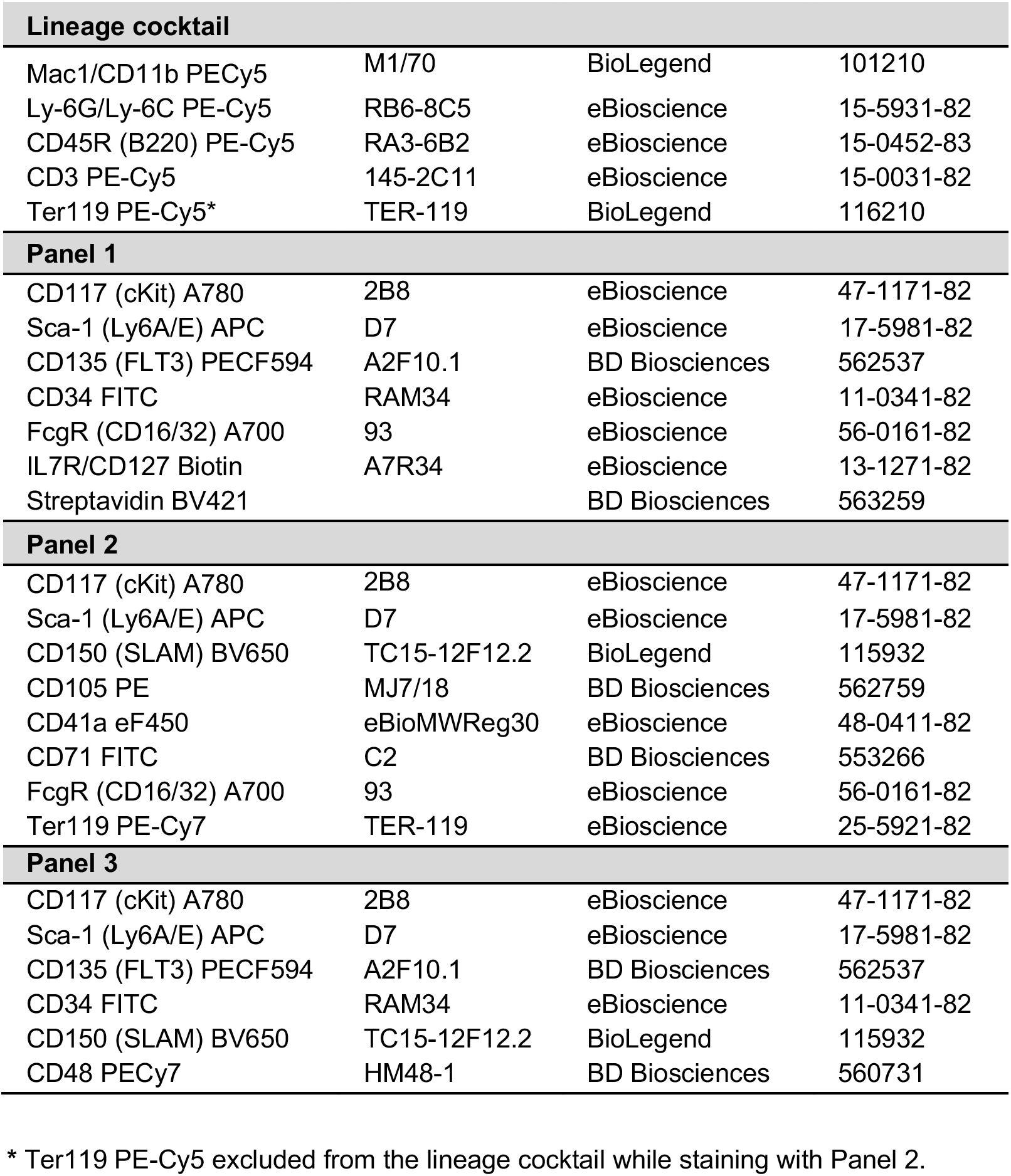

| Antibody | Clone | Source | Cat number |
| --- | --- | --- | --- |

| **Lineage cocktail** |  |  |  |
| --- | --- | --- | --- |
| Mac1/CD11b PECy5 | M1/70 | BioLegend | 101210 |
| Ly-6G/Ly-6C PE-Cy5 | RB6-8C5 | eBioscience | 15-5931-82 |
| CD45R (B220) PE-Cy5 | RA3-6B2 | eBioscience | 15-0452-83 |
| CD3 PE-Cy5 | 145-2C11 | eBioscience | 15-0031-82 |
| Ter119 PE-Cy5* | TER-119 | BioLegend | 116210 |
| **Panel 1** |  |  |  |
| CD117 (cKit) A780 | 2B8 | eBioscience | 47-1171-82 |
| Sca-1 (Ly6A/E) APC | D7 | eBioscience | 17-5981-82 |
| CD135 (FLT3) PECF594 | A2F10.1 | BD Biosciences | 562537 |
| CD34 FITC | RAM34 | eBioscience | 11-0341-82 |
| FcgR (CD16/32) A700 | 93 | eBioscience | 56-0161-82 |
| IL7R/CD127 Biotin | A7R34 | eBioscience | 13-1271-82 |
| Streptavidin BV421 |  | BD Biosciences | 563259 |
| **Panel 2** |  |  |  |
| CD117 (cKit) A780 | 2B8 | eBioscience | 47-1171-82 |
| Sca-1 (Ly6A/E) APC | D7 | eBioscience | 17-5981-82 |
| CD150 (SLAM) BV650 | TC15-12F12.2 | BioLegend | 115932 |
| CD105 PE | MJ7/18 | BD Biosciences | 562759 |
| CD41a eF450 | eBioMWReg30 | eBioscience | 48-0411-82 |
| CD71 FITC | C2 | BD Biosciences | 553266 |
| FcgR (CD16/32) A700 | 93 | eBioscience | 56-0161-82 |
| Ter119 PE-Cy7 | TER-119 | eBioscience | 25-5921-82 |
| **Panel 3** |  |  |  |
| CD117 (cKit) A780 | 2B8 | eBioscience | 47-1171-82 |
| Sca-1 (Ly6A/E) APC | D7 | eBioscience | 17-5981-82 |
| CD135 (FLT3) PECF594 | A2F10.1 | BD Biosciences | 562537 |
| CD34 FITC | RAM34 | eBioscience | 11-0341-82 |
| CD150 (SLAM) BV650 | TC15-12F12.2 | BioLegend | 115932 |
| CD48 PECy7 | HM48-1 | BD Biosciences | 560731 |

\* Ter119 PE-Cy5 excluded from the lineage cocktail while staining with Panel 2.

### Evotip Pure protocol for 500 cells mouse BM samples

The 384-well plates were thawed in the fridge and each well was loaded onto Evotip Pure according to the manufacturer’s instructions. Briefly, the tips were rinsed with 20 μl Solvent B (MS-grade ACN with 0.1% formic acid), conditioned by soaking in 2-propanol for 30 sec, equilibrated with 20 μl Solvent A (MS-grade water with 0.1% formic acid). The centrifugation steps in between were always at 700 g for 60 seconds. The tips were placed in a box filled with Solvent A to keep Evotips wet while sample loading.

Each well containing 500 cells (around 4.5 µl) was loaded on wet tips containing 15 μl solvent A, the wells were rinsed with 5 µl Buffer A* (2% ACN, 0.1% TFA) and loaded onto tips. The loaded tips were centrifuged at 700 g for 60 seconds and washed with 20 μl solvent A. Finally, tips were filled with 250 μl Solvent A, and submerged in a box with Solvent A.

### Liquid chromatography configuration

Chromatographic separation of peptides was conducted on an EvosepOne UHPLC system (Evosep) connected to a 15 cm Aurora Elite™ TS (Ion Opticks). All separations were carried out with the column oven set to 50°C and utilizing Evosep’s built-in 20 samples per day (SPD) method.

### Mass spectrometry data acquisition

The acquisition of peptides from 500 cells was conducted using a Thermo Scientific Orbitrap Eclipse Tribrid mass spectrometer, which was operated in positive mode and equipped with the FAIMS Pro interface. For this process, a single compensation voltage of −45 V was utilized, as detailed in the methodology of Petrosius et al., 2023 ^17^. The MS1 spectra were obtained using the Orbitrap, with a set resolution of 120,000 and a scan range between 400 and 1000 m/z. Additionally, the normalized automatic gain control (AGC) target was set at 300%, alongside a maximum injection time of 246 ms.

Data-independent acquisition for MS2 spectra was carried out in the Orbitrap at the same resolution of 120,000. Loop control was set to 12 spectra per loop, utilizing isolation windows of 17 Th and a mass range spanning from 200 to 1200 Th. This resulted in a total of 36 windows across all looped cycles. A 1 m/z window overlap was employed, and the fragmentation of precursor ions was achieved through higher energy collisional dissociation (HCD). The normalized collision energy (NCE) used for fragmentation was set to 33%. The AGC target for this was set at 1000%, and the maximum injection time was configured to automatic.

### Data analysis

The raw data was processed with Spectronaut in directDIA mode with standard setting with the following modifications: Quantity MS level was changed to MS1 and Carbamidomethylation of cysteines was removed as fixed modification. Protein quantification matrices were then exported and further analysis with custom python and R scripts with Visual Studio Code (1.84.2).

### Transcription factor clustering

Transcription factors were clustered based on their abundances in each population. Missing values were set to zero, to avoid filtering out transcription factors that are expressed only in specific populations. A one-way ANOVA test was carried out upstream of clustering to filter out proteins that are not significantly altered between all the groups (Qvalue 0.001 cut-off). An affinity propagation clustering approach was used which finds exemplar cases that can be best used to represent the complete dataset. The algorithm was implemented via the sklearn python package, the default damping parameter of 0.5 was used and the number of clustered set to the median of the input similarities. Proteins were annotated as transcription factors based on the AnimalTFDB v4.0.

### Cell surface marker inference

The list of potential cell surface proteins was composed by concatenative the cell surface protein catalog from BD and the Cell surface protein atlas (CSPA) ^26^. The CSPA was further filtered for only proteins that were categorized as “high confidence” and had at least one predicted transmembrane domain. The concatenated list was then cross-referenced with the proteomics data. Two approaches were chosen to identify the most promising cell surface proteins, first limma was used to apply a linear to model to calculate the log2FC and statistical significance of the different contrast between the target population and the background cKit population (for HSC CD150+ and MPP CD150-LSK was used). The mutual information values were calculated for each cell surface proteins for a given population to obtain a second metric, which is routinely used in decision tree based models. The top proteins were then selected and visualized further to showcase how well they could separate populations of interest.

## Data availability

The data matrices are available with https://doi.org/10.5281/zenodo.10417802

## Supplementary Figures

**Figure S1.**
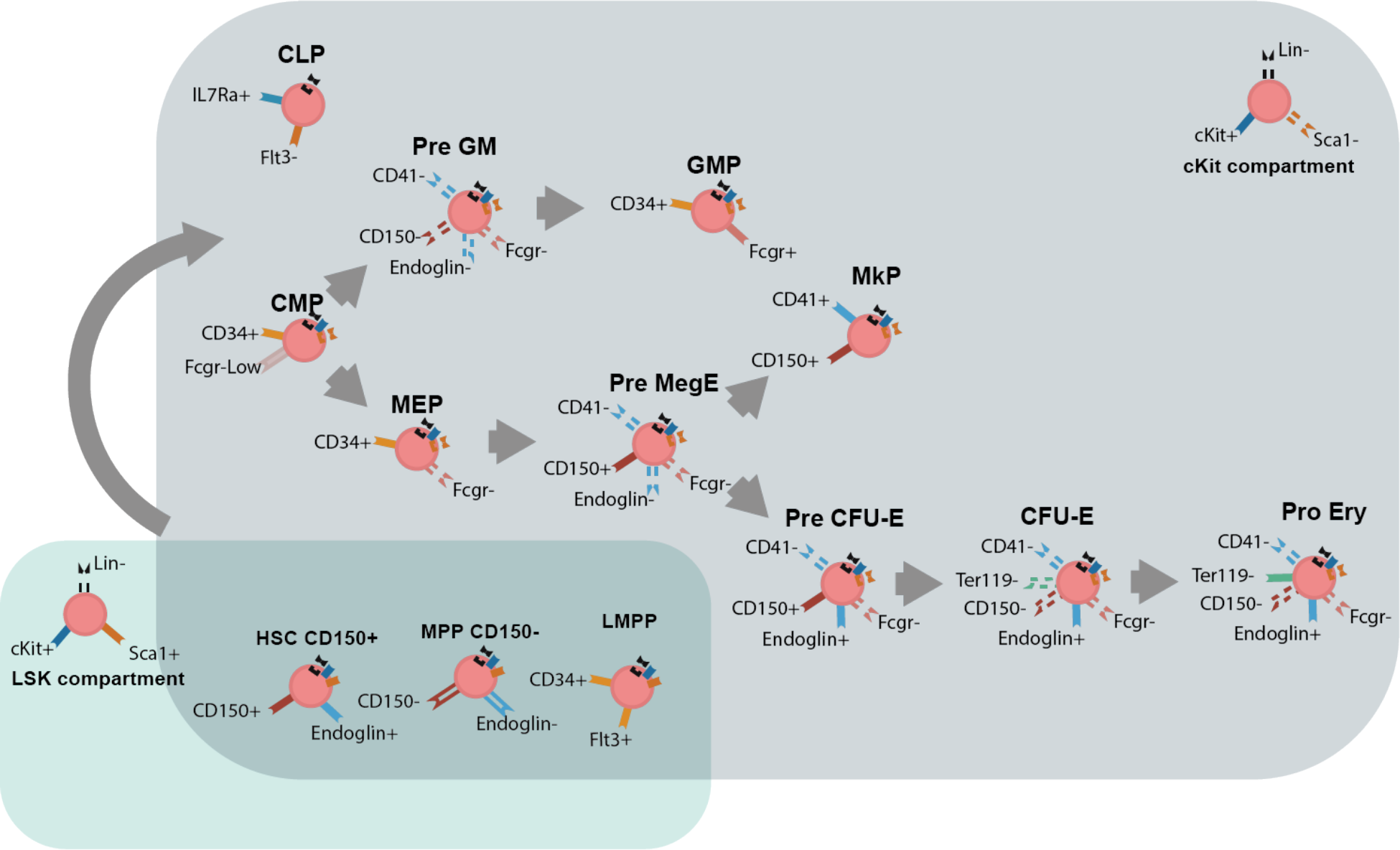
Illustration of the isolated hematopoietic populations. The color blocks denote two compartments LSK and cKit, the FACS markers are noted with stick figures, where full figure indicates positive and dashed empty negative selection. Only the markers noted on the cells are used for isolation.

**Figure S2.**
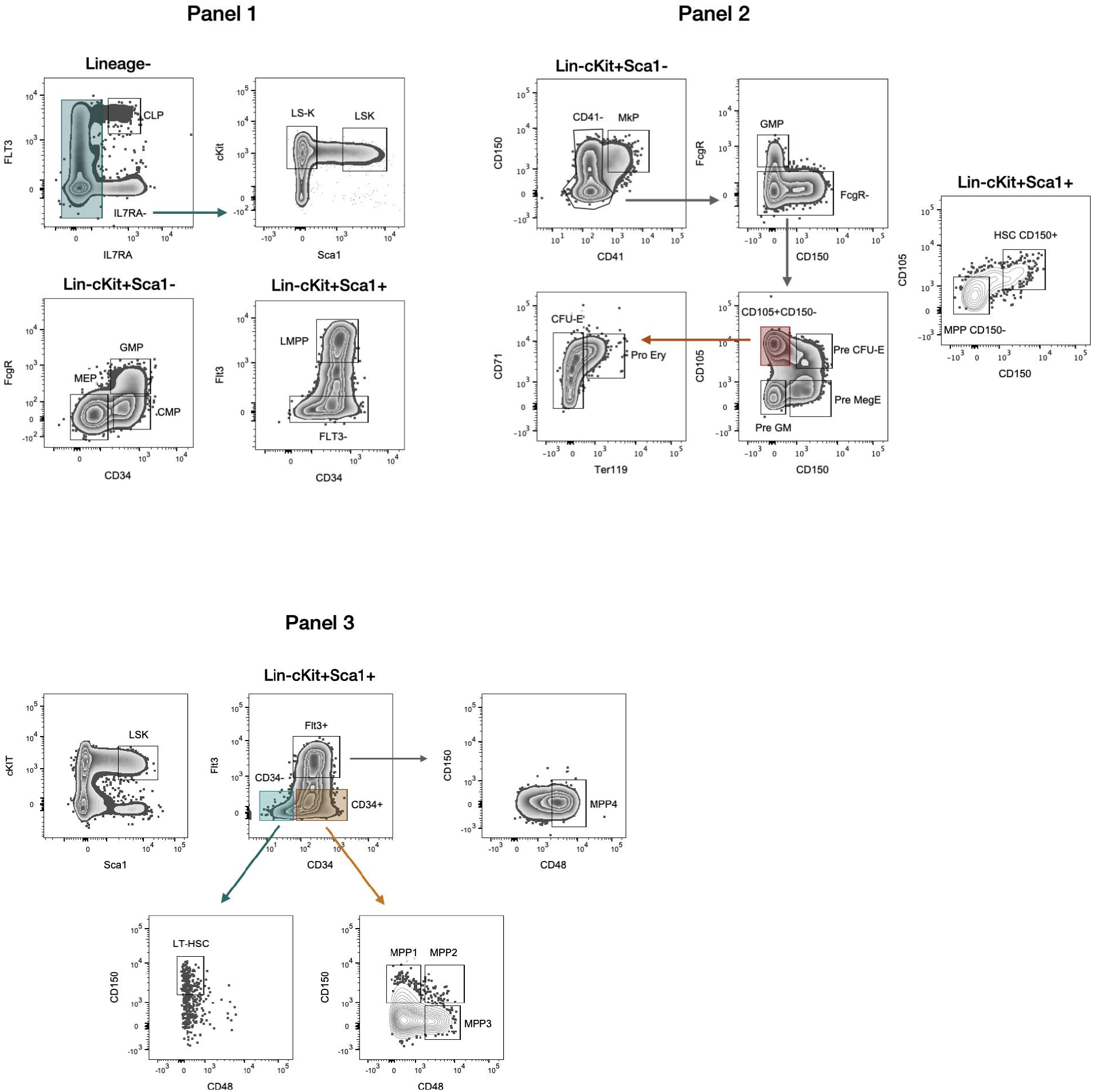
FACS gating strategies employed in the low-input proteomics dataset. The panels for the “one mouse hematopoietic hierarchy” (Panel 1 and Panel 2) were designed based on the gating strategy outlined in Akashi K. et al. 2000 and Pronk C. et al. 2007. For the early stem cell compartment, the panel 3 was designed following the gating strategy outlined in Cabezas-Wallscheid et al. 2014.

**Figure S3.**
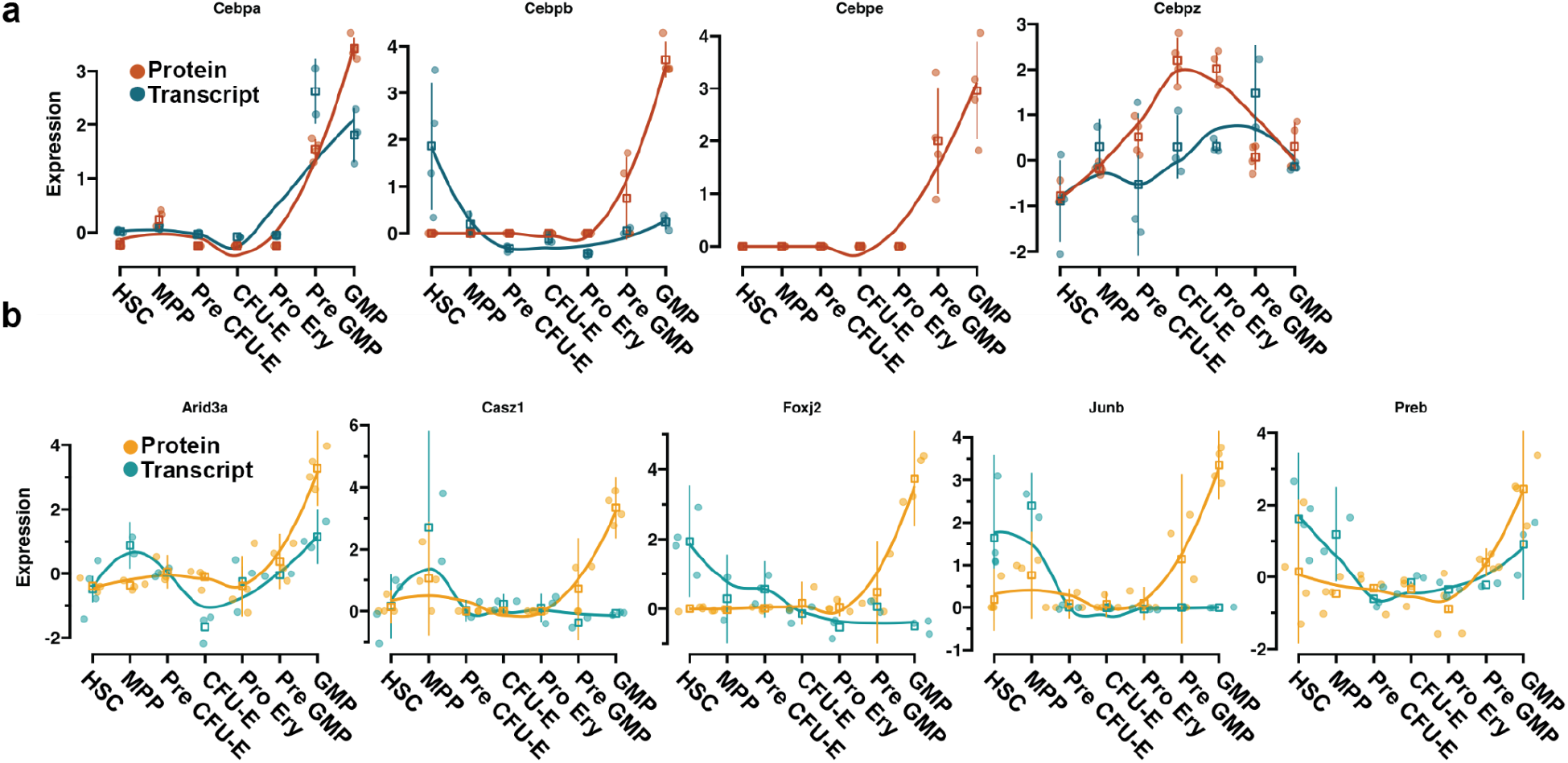
Comparison of GMP specific transcription factor protein and transcript levels in different selected populations. a) Comparison of known CEBP protein levels on transcript (blue) and protein level (red). b) Protein and transcript expression of tentative GMP specific transcription factors. Round dots note individual measurement values, square the mean and the line represent the standard deviation. The transcriptome level data is used from Pronk et al, 2007 ^12^, that used the same population isolation scheme. The values were downloaded via the BloodSpot web application.

**Figure S4.**
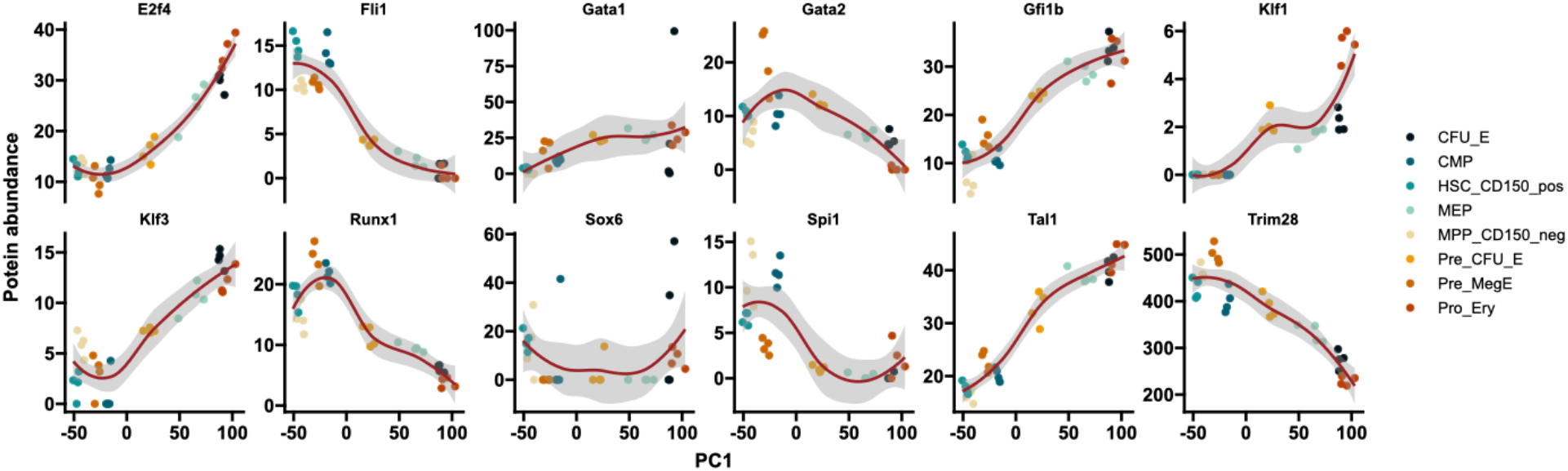
Comparison of transcription factors showcases in Gillespie et al 2020 ^13^. Scatter plots with fitted smooth line showing the trends of different transcription factors showcased in previous study. The first principal component (PC1) was used as the x-axis as it reflected the erythroid differentiation trajectory. Y-axis represents the log2 transformed protein abundance and the point color specific populations. Loess is used to fit the trend line and the grey area notes the confidence interval.

## Full clusters from Figure 2

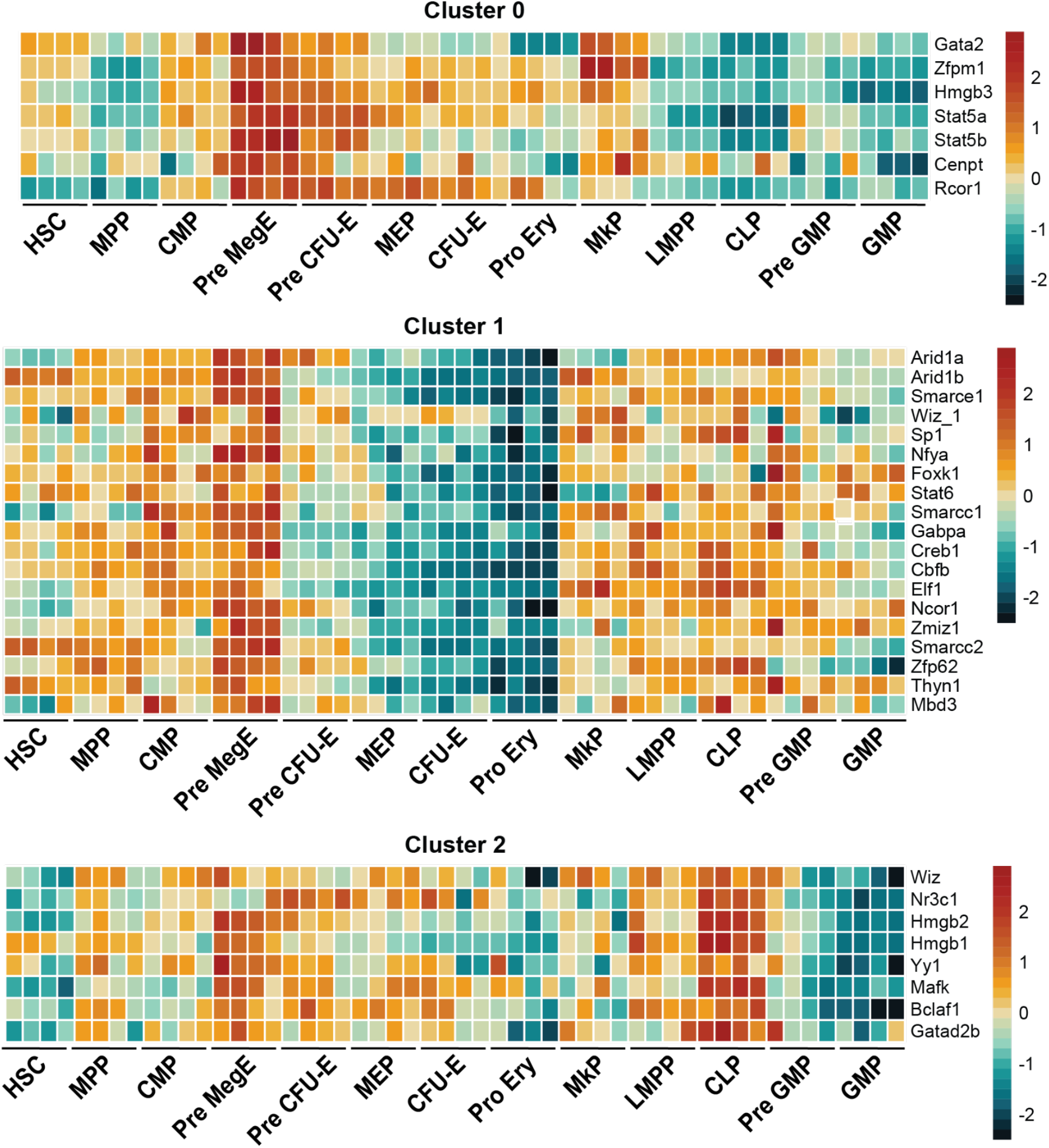
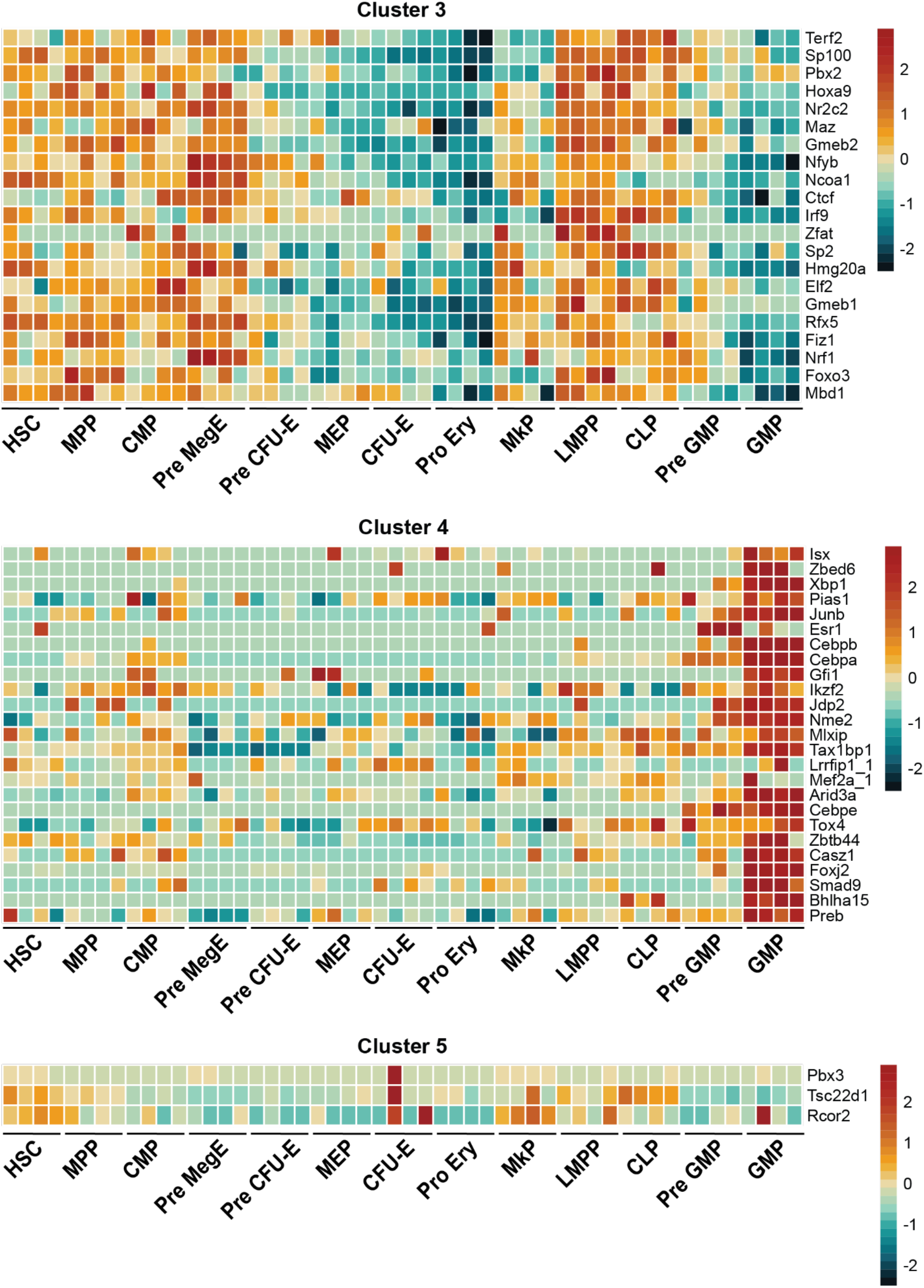
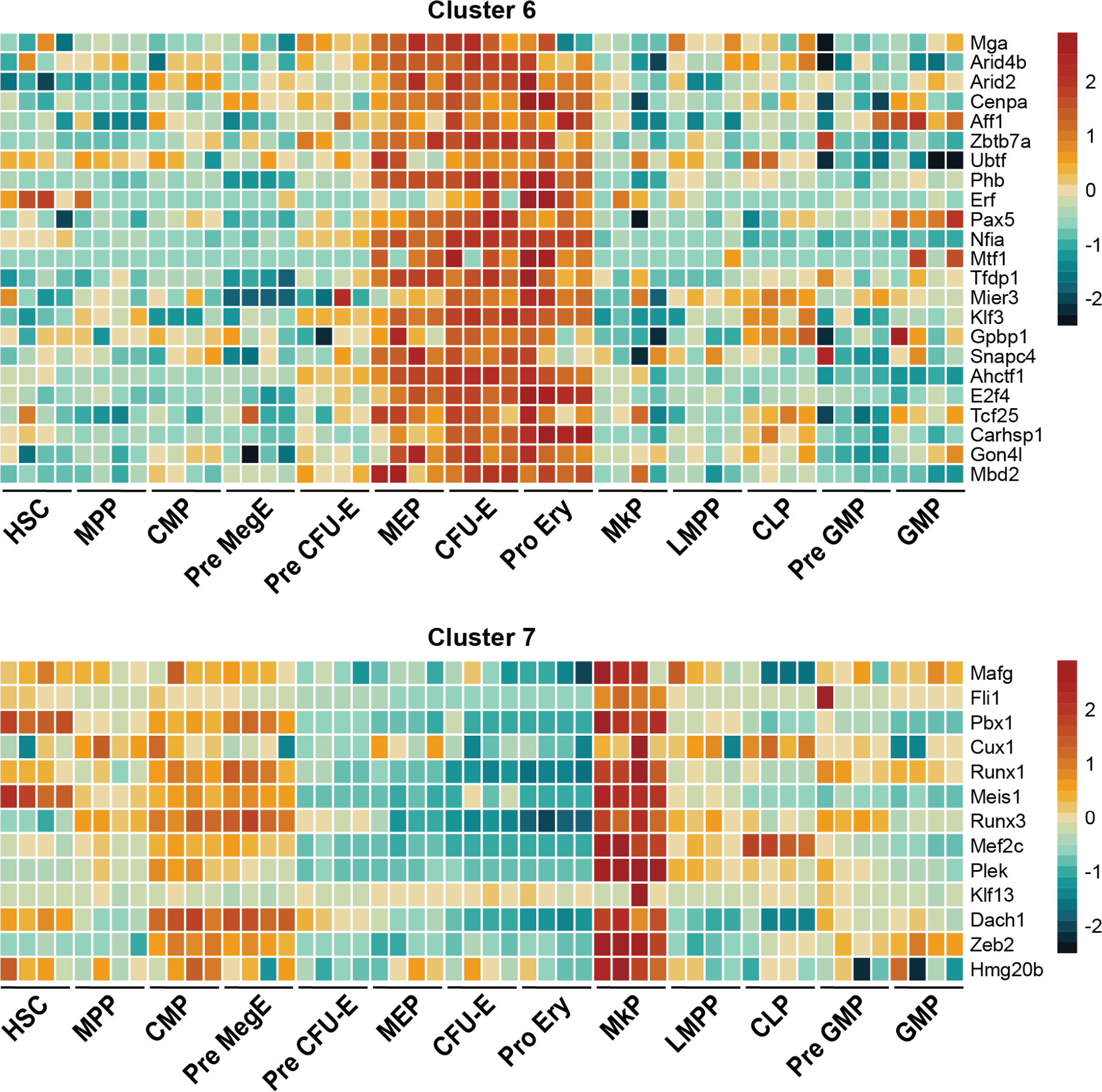
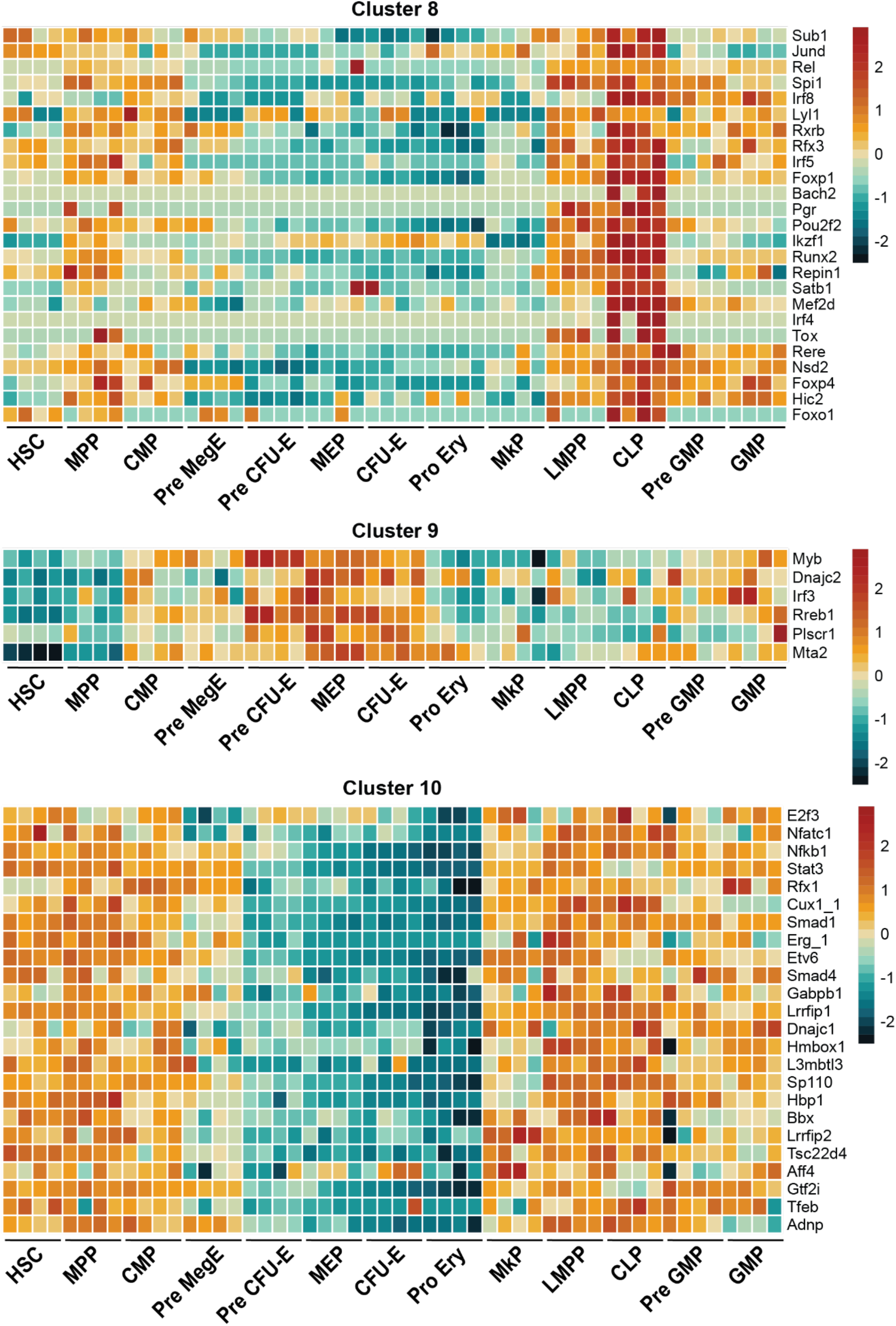
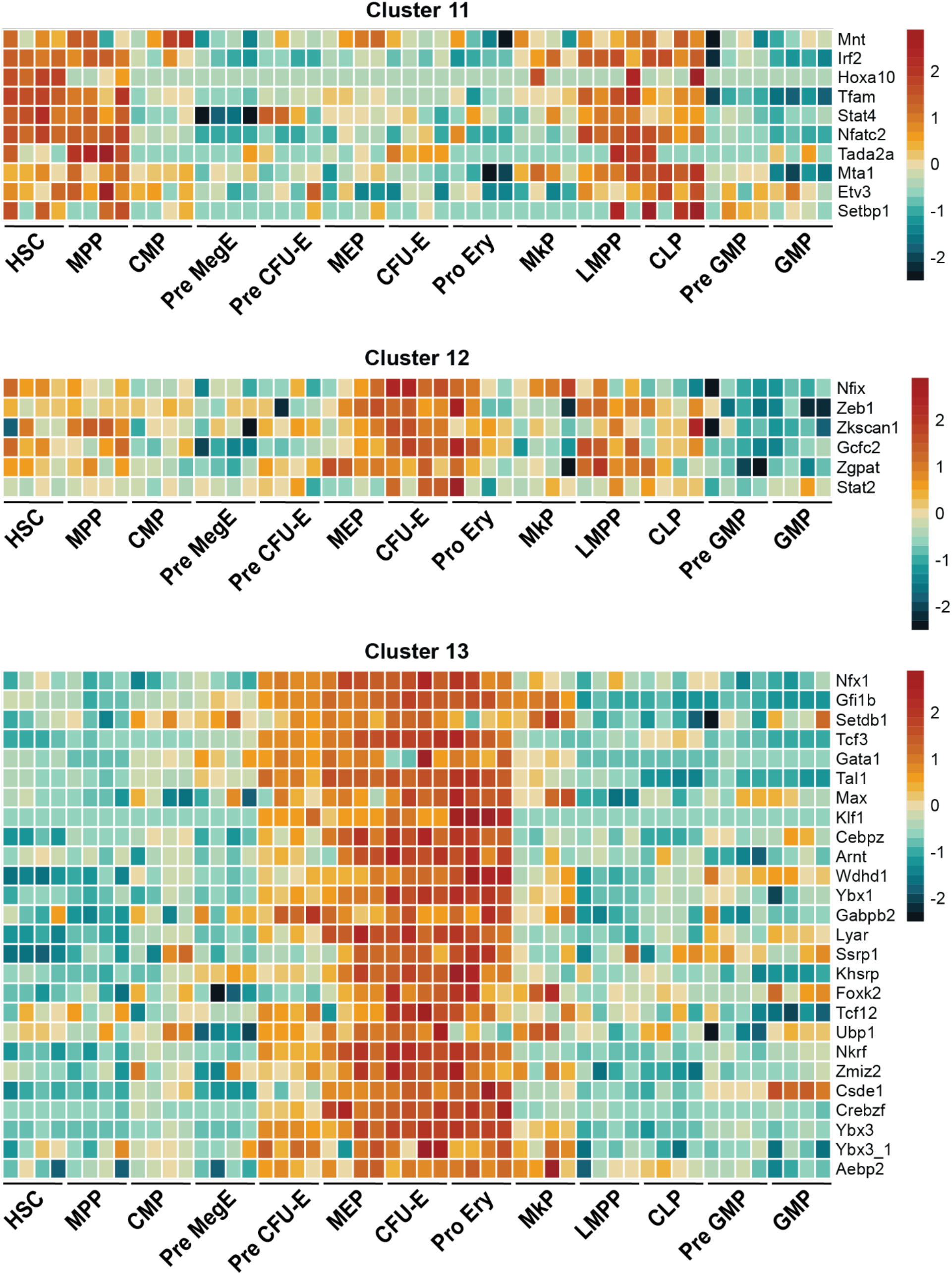
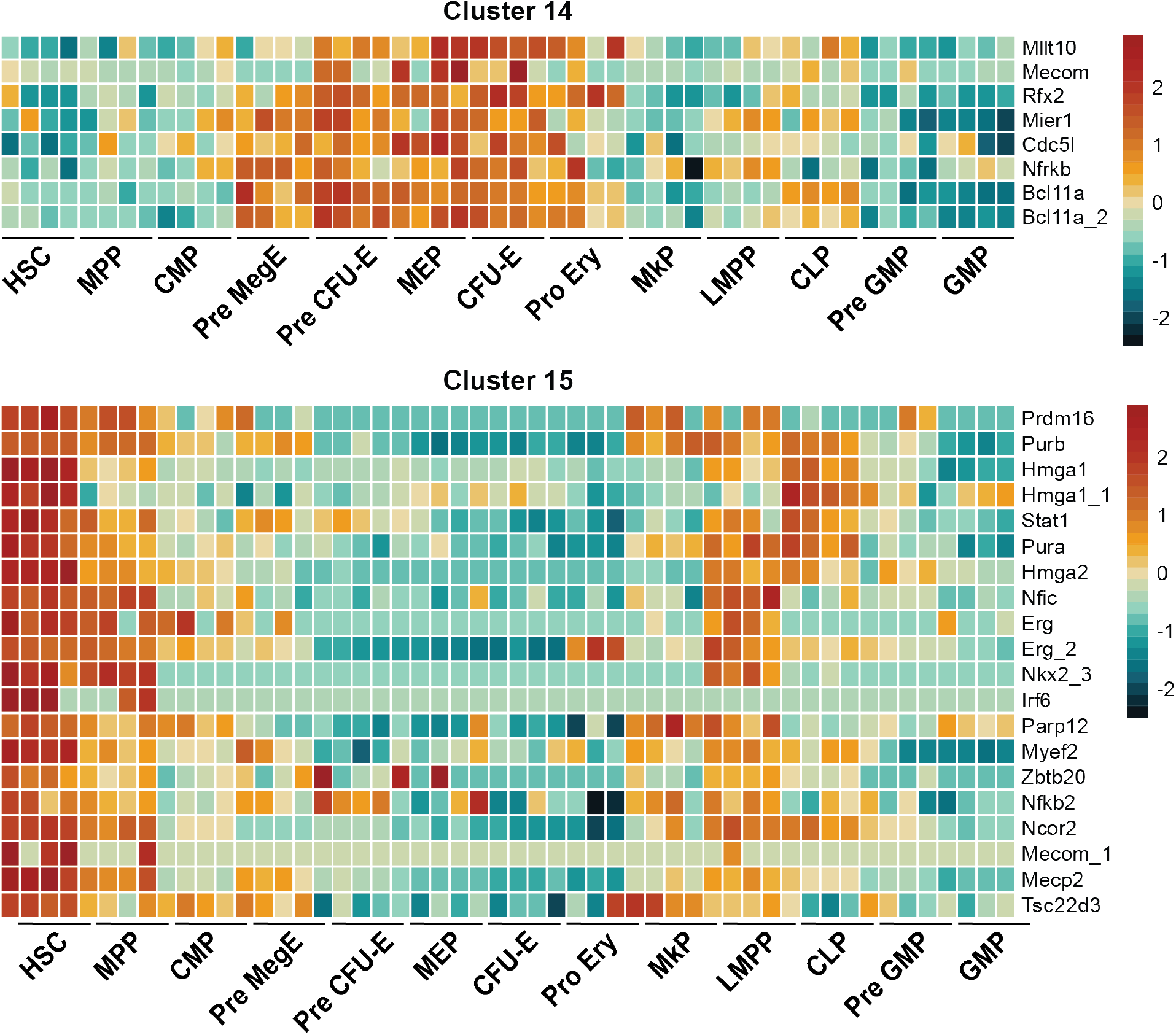

## Author contributions

B.T.P., E.M.S., N.Ü. and V.P. conceived and designed the project. N.Ü. and P.A.F. performed experiments. P.A.F conducted the MS data acquisition. V.P. performed the data analysis and data visualization. B.F. provided critical input. The manuscript was drafted and revised by N.Ü. and V.P. with input from E.M.S. and B.T.P. The work was supervised by B.T.P. and E.M.S.

## Acknowledgments

Work in the Porse lab was supported by grants from the Candys Foundation, the Danish Cancer Society, the Eva and Henrik Frænkels memorial Foundation and through a center grant from the Novo Nordisk Foundation (Novo Nordisk Foundation Center for Stem Cell Biology, DanStem; Grant Number NNF17CC0027852). Work in the Schoof lab was supported by grants from the Independent Research Fund Denmark (2067-00053B), Danish Cancer Society (R324-A17978), Novo Nordisk Foundation (NNF21OC0071016), Leo Foundation (LF-OC-21-000832) and the Lundbeck Foundation (R413-2022-869).

## References

1. Orkin, S. H. & Zon, L. I. Hematopoiesis: An Evolving Paradigm for Stem *Cell* Biology. Cell 132, 631–644 (2008).

2. Ikuta, K. & Weissman, I. L. Evidence that hematopoietic stem cells express mouse c-kit but do not depend on steel factor for their generation. Proc. Natl. Acad. Sci. 89, 1502– 1506 (1992).

3. Akashi, K., Traver, D., Miyamoto, T. & Weissman, I. L. A clonogenic common myeloid progenitor that gives rise to all myeloid lineages. Nature 404, 193–197 (2000).

4. Kondo, M., Weissman, I. L. & Akashi, K. Identification of Clonogenic Common Lymphoid Progenitors in Mouse Bone Marrow. Cell 91, 661–672 (1997).

5. Seita, J. & Weissman, I. L. Hematopoietic stem cell: self-renewal versus differentiation. Wiley Interdiscip. Rev.: Syst. Biol. Med. 2, 640–653 (2010).

6. Osawa, M., Hanada, K., Hamada, H. & Nakauchi, H. Long-Term Lymphohematopoietic Reconstitution by a Single CD34-Low/Negative Hematopoietic Stem Cell. Science 273, 242–245 (1996).

7. Kiel, M. J. et al. SLAM Family Receptors Distinguish Hematopoietic Stem and Progenitor *Cells* and Reveal Endothelial Niches for Stem Cells. Cell 121, 1109–1121 (2005).

8. Wilson, A. et al. Hematopoietic Stem Cells Reversibly Switch from Dormancy to Self-Renewal during Homeostasis and Repair. Cell 135, 1118–1129 (2008).

9. Cabezas-Wallscheid, N. et al. Identification of Regulatory Networks in HSCs and Their Immediate Progeny via Integrated Proteome, Transcriptome, and DNA Methylome Analysis. Cell Stem Cell 15, 507–522 (2014).

10. Pietras, E. M. et al. Functionally Distinct Subsets of Lineage-Biased Multipotent Progenitors Control Blood Production in Normal and Regenerative Conditions. Cell Stem Cell 17, 35–46 (2015).

11. Rodriguez-Fraticelli, A. E. et al. Clonal analysis of lineage fate in native haematopoiesis. Nature 553, 212–216 (2018).

12. Pronk, C. J. H. et al. Elucidation of the Phenotypic, Functional, and Molecular Topography of a Myeloerythroid Progenitor Cell Hierarchy. Cell Stem Cell 1, 428–442 (2007).

13. Gillespie, M. A. et al. Absolute Quantification of Transcription Factors Reveals Principles of Gene Regulation in Erythropoiesis. Mol Cell 78, 960–974.e11 (2020).

14. Signer, R. A. J., Magee, J. A., Salic, A. & Morrison, S. J. Haematopoietic stem cells require a highly regulated protein synthesis rate. Nature 509, 49–54 (2014).

15. Zaro, B. W. et al. Proteomic analysis of young and old mouse hematopoietic stem cells and their progenitors reveals post-transcriptional regulation in stem cells. Elife 9, e62210 (2020).

16. Amon, S. et al. Sensitive Quantitative Proteomics of Human Hematopoietic Stem and Progenitor Cells by Data-independent Acquisition Mass Spectrometry* [S]. Mol Cell Proteomics 18, 1454–1467 (2019).

17. Petrosius, V. et al. Exploration of cell state heterogeneity using single-cell proteomics through sensitivity-tailored data-independent acquisition. Nat. Commun. 14, 5910 (2023).

18. Graf, T. & Enver, T. Forcing cells to change lineages. Nature 462, 587–594 (2009).

19. Laiosa, C. V., Stadtfeld, M., Xie, H., de Andres-Aguayo, L. & Graf, T. Reprogramming of Committed T Cell Progenitors to Macrophages and Dendritic Cells by C/EBPα and PU.1 Transcription Factors. Immunity 25, 731–744 (2006).

20. Rosenbauer, F. & Tenen, D. G. Transcription factors in myeloid development: balancing differentiation with transformation. Nat. Rev. Immunol. 7, 105–117 (2007).

21. Cantor, A. B. & Orkin, S. H. Transcriptional regulation of erythropoiesis: an affair involving multiple partners. Oncogene 21, 3368–3376 (2002).

22. Iwasaki, H. et al. GATA-1 Converts Lymphoid and Myelomonocytic Progenitors into the Megakaryocyte/Erythrocyte Lineages. Immunity 19, 451–462 (2003).

23. Copley, M. R. et al. The Lin28b–let-7–Hmga2 axis determines the higher self-renewal potential of fetal haematopoietic stem cells. Nat. Cell Biol. 15, 916–925 (2013).

24. Passegué, E., Jochum, W., Schorpp-Kistner, M., Möhle-Steinlein, U. & Wagner, E. F. Chronic Myeloid Leukemia with Increased Granulocyte Progenitors in Mice Lacking JunB Expression in the Myeloid Lineage. Cell 104, 21–32 (2001).

25. Bagger, F. O. et al. BloodSpot: a database of gene expression profiles and transcriptional programs for healthy and malignant haematopoiesis. Nucleic Acids Res. 44, D917–D924 (2016).

26. Bausch-Fluck, D. et al. A Mass Spectrometric-Derived Cell Surface Protein Atlas. Plos One 10, e0121314 (2015).

27. Bausch-Fluck, D. et al. The in silico human surfaceome. Proc National Acad Sci 115, E10988–E10997 (2018).

28. Hopp, A.-K., Grüter, P. & Hottiger, M. O. Regulation of Glucose Metabolism by NAD+ and ADP-Ribosylation. Cells 8, 890 (2019).

29. Wilson, A. & Trumpp, A. Bone-marrow haematopoietic-stem-cell niches. Nat. Rev. Immunol. 6, 93–106 (2006).

30. Bianchi, M. E. & Agresti, A. HMG proteins: dynamic players in gene regulation and differentiation. Curr. Opin. Genet. Dev. 15, 496–506 (2005).

31. Pfannkuche, K., Summer, H., Li, O., Hescheler, J. & Dröge, P. The High Mobility Group Protein HMGA2: A Co-Regulator of Chromatin Structure and Pluripotency in Stem Cells? Stem Cell Rev. Rep. 5, 224–230 (2009).

32. Minervini, A. et al. HMGA Proteins in Hematological Malignancies. Cancers 12, 1456 (2020).

33. Yuan, S., Liu, Z., Xu, Z., Liu, J. & Zhang, J. High mobility group box 1 (HMGB1): a pivotal regulator of hematopoietic malignancies. J. Hematol. Oncol. 13, 91 (2020).

34. Morishita, A. et al. HMGA2 Is a Driver of Tumor Metastasis. Cancer Res. 73, 4289–4299 (2013).

35. West, A. P. & Shadel, G. S. Mitochondrial DNA in innate immune responses and inflammatory pathology. Nat. Rev. Immunol. 17, 363–375 (2017).

36. Amir, R. E. et al. Rett syndrome is caused by mutations in X-linked MECP2, encoding methyl-CpG-binding protein 2. Nat. Genet. 23, 185–188 (1999).

37. Simicevic, J. et al. Absolute quantification of transcription factors during cellular differentiation using multiplexed targeted proteomics. Nat. Methods 10, 570–576 (2013).

38. Matzinger, M., Müller, E., Dürnberger, G., Pichler, P. & Mechtler, K. Robust and Easy-to-Use One-Pot Workflow for Label-Free Single-Cell Proteomics. Anal. Chem. 95, 4435– 4445 (2023).

39. Schoof, E. M. et al. Quantitative single-cell proteomics as a tool to characterize cellular hierarchies. Nat Commun 12, 3341 (2021).

